# Mental Effort Cost Learning is Retrospective

**DOI:** 10.1101/2022.11.26.518013

**Authors:** Asako Mitsuto, Rei Akaishi, Keiichi Onoda, Kenji Morita, Toshikazu Kawagoe, Tetsuya Yamamoto, Shuhei Yamaguchi, Ritsuko Hanajima, Andrew Westbrook

**Affiliations:** Department of Psychiatry, Center for Advanced Human Brain Imaging Research, Rutgers University, Piscataway, NJ 08854, USA; Division of Neurology, Department of Brain and Neurosciences, Faculty of Medicine, Tottori University, 86-1 Nishi-Cho, Yonago 683-8504, Japan; Research Fellowship for Young Scientists, Japan Society for the Promotion of Science, Tokyo 102-0083, Japan; Physical and Health Education, Graduate School of Education, The University of Tokyo, Hongo 7-3-1, Bunkyo-Ku, Tokyo 113-0033, Japan; Department of Neurology, Faculty of Medicine, Shimane University, Enya-Cho 89-1, Izumo 693-8501, Japan; Department of Cognitive Neuroscience, Graduate School of Medicine, The University of Tokyo, 7-3-1 Hongo, Bunkyo-Ku, Tokyo 113-0033, Japan; Center for Brain Science, RIKEN, Wako, Saitama 351-0198, Japan; Faculty of Psychology, Otemon Gakuin University, Ibaraki, Osaka 567-8502, Japan; International Research Center for Neurointelligence (WPI-IRCN), The University of Tokyo, 7-3-1 Hongo, Bunkyo-ku, Tokyo 113-0033, Japan; School of Humanities and Science, Kumamoto Campus, Tokai University, Toroku 9-1-1, Kumamoto-city, Kumamoto 862-8652, Japan; Brain Function Imaging and Support Center, Laboratory of Biofunctional Information Analysis, National Institute for Physiological Sciences; Department of Neurology, Shimane Prefectural Central Hospital, Izumo 693-8555, Japan

**Author notes:** These senior authors contributed equally (Shuhei Yamaguchi, Ritsuko Hanajima, Andrew Westbrook).

**Keywords:** frontal cortex, striatum, mental effort, reinforcement learning, fMRI, prediction error, uncertainty, temporal-difference learning

## Abstract

To understand why people avoid mental effort, it is crucial to reveal the mechanisms by which we learn and decide about mental effort costs. This study investigated whether mental effort cost learning aligns with temporal-difference (TD) learning or alternative mechanisms. Model-based fMRI analyses showed no correlation between cost prediction errors (CPEs) and activity in the dorsomedial frontal cortex/dorsal anterior cingulate cortex (dmFC/dACC) or striatum at the time of a fully informative effort cue about upcoming effort demands, contradicting the TD hypothesis. Instead, CPEs correlate with dmFC/dACC (positively) and caudate (negatively) activity at effort completion. Furthermore, only activity patterns at effort completion predict subsequent choices. These results show that decision policies are updated retrospectively at effort completion, updating expected costs with prediction error between experienced effort and prior expectations, demonstrating mental effort cost learning is retrospective, and imply that adaptive learning of mental effort cost does not follow canonical TD learning.

**Significance Statement:** Understanding how people learn about mental effort costs is essential for advancing theories of motivation and cognitive control. However, the algorithms supporting such learning remain unclear. This study addressed this gap and found that temporal-difference learning, commonly used to explain reward learning, could not account for how people learn about effort. Instead, decision policies were updated retrospectively at effort completion, based on a prediction error between experienced effort and prior expectations. These findings reveal that mental effort cost learning is fundamentally retrospective and imply that it relies on mechanisms distinct from canonical temporal-difference learning.

## Introduction

People exert effort adaptively to achieve their goals. Understanding goal-directed behavior requires revealing the mechanisms by which people learn and decide about effort. To make effort-based decisions, humans and animals integrate effort costs and benefits. Reward learning mechanisms are well understood and effort cost-benefit evaluation has been studied extensively^1–8^. However, the neural mechanisms by which mental effort costs are learned remains unclear^9–15^.

A central question is what algorithms govern mental effort cost learning. Like rewards, effort may be learned through reinforcement learning (RL), with the brain generating cost prediction errors (CPEs)^11–13^. However, when and how CPEs are generated remains unknown. Negatively-valenced stimuli such as pain, air puff, or reward omission are learned via temporal-difference (TD) learning^16–18^. TD algorithms involve an update of the estimated value of future outcomes, based on the difference between value estimates immediately before and after information about reinforcers becomes available. Thus, in TD learning, informative cues should evoke a CPE signal. However, evidence suggests that physical or mental effort demands appear to be encoded after effort is exerted, or even retrospectively, at the time of subsequent reward^19–21^. Because prior studies have not dissociated effort production from effort information, it remains unclear whether information about mental effort demands alone is sufficient to trigger CPE signaling, as predicted by TD learning. Moreover, because effort encoding has mostly been studied in terms of reward discounting^20–24^, it is unclear how effort cost learning proceeds in the absence of reward processing. In this study, we use fMRI to examine neural dynamics while humans learn about mental effort costs, in the absence of rewards, in a task where information about upcoming effort demands is separated in time from effort exertion itself.

Initially, we hypothesized that mental effort cost learning, like reward learning, is governed by TD learning. Thus, we predicted that informative cues about upcoming effort demands would trigger CPE signaling. Moreover, we predicted that CPEs would, at the time of effort cues, correlate positively with the activity in the dmFC/dACC, and negatively with the activity in the striatum. Our predictions were based on three lines of evidence. First, monkey electrophysiology data reveals that the firing rates of distinct populations of mPFC neurons encode the PE of rewards and punishments positively^25,26^. Second, our previous fMRI study for humans showed that the activity in the dmFC/dACC and other regions correlates positively with mental effort CPE signals^11^ – though in that study, we did not dissociate effort exertion from effort information processing. Third, human fMRI data reveal a positive correlation of PE of physical effort with the activity in the dmFC^24^.

There is also reason to suspect that CPE signals correlate negatively with activity in the striatum. In terms of anatomy, parallel cortico-basal ganglia loops are a likely substrate for effort cost learning given their role in action selection via dopamine-mediated plasticity^4,27,28^. If reward PEs reflect integrated utility, CPEs should correlate negatively with striatal activity, contrasting with a positive correlation for reward PEs^29–31^. We note, however, that PEs for positive punishment, such as pain and loss of financial reward, correlate positively with striatal activity^17,29^. As such, the sign of CPE encoding in the dmFC/dACC and the striatum at the time of the effort cues is uncertain.

To test our hypotheses, we designed two fMRI experiments of demand-selection tasks with informative cues about upcoming effort which would, according to TD learning, permit participants to generate CPE signals based on the differences between expected effort ^11^costs before and after effort cues. In a trial, participants chose between two options associated probabilistically with either high or low demand mental arithmetic and spatial reasoning tasks (Figure 1A). High/low demand probabilities for each option changed over time. We next presented effort cues which were fully informative of upcoming effort levels. The first hypothesis predicted that, at the time of effort cues, brains represent CPE signals (Figure 4Aa). Contrary to the first hypothesis, CPE signals did not correlate with the activity in the dmFC/dACC or striatum, at the time of effort cues. We thus speculated that mental effort cost learning does not follow canonical TD learning and considered alternative hypotheses. Specifically, we hypothesized that expected costs may instead be updated at the point of effort initiation, when participants have committed to an action (Figure 4Ab). Although we found that CPE signals correlated positively with the activity of the ventromedial prefrontal cortex (vmPFC)/ventral anterior cingulate cortex (vACC)/orbitofrontal cortex (OFC), this result did not confirm our second hypothesis about the anatomical location of the CPE which we predicted would involve the anterior cingulate cortex and the striatum. We next hypothesized that an update of expected effort requires effort completion, and thus we predicted that the brain would encode CPEs at the moment that effort is concluded (Figure 4Ac). Confirming our third hypothesis, CPE signals correlated positively with the activity of the dmFC/dACC and other regions, and negatively in the bilateral caudate nucleus (Cd), at the time of effort completion. Moreover, a test of the timing of choice policy updating revealed that residual blood oxygenation level dependent (BOLD) signals predicted the next trials’ choice (stay/switch) only at the time of effort completion. These results suggest that the adaptive learning of mental effort cost is retrospective.

**Figure 1.**
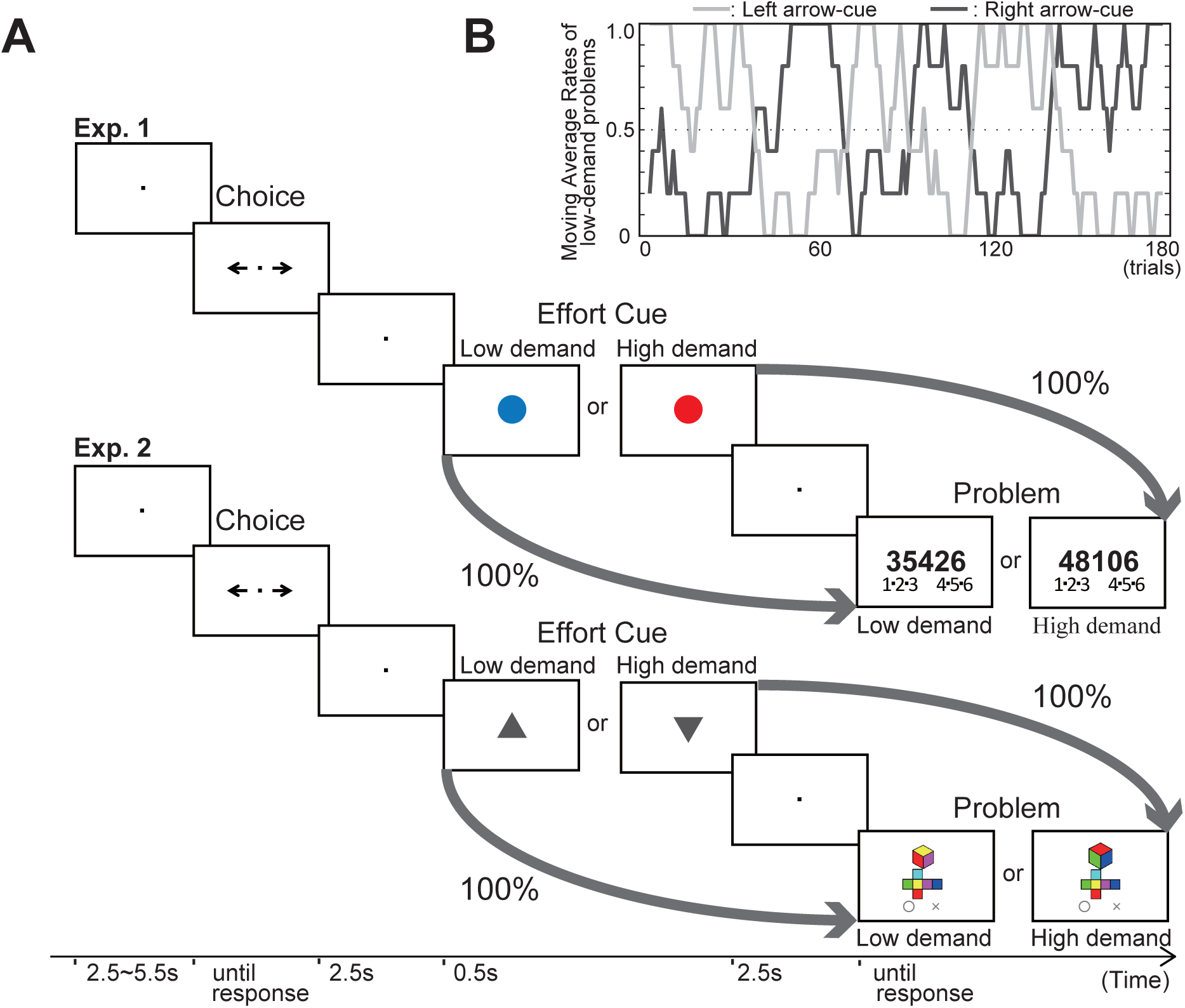
The behavioral paradigm of demand-selection tasks with effort cues. (A) At the beginning of each trial, participants selected either a left or right arrow associated probabilistically with either low- or high-demand problems. After an effort cue, a (low- or high-demand) problem was presented. Effort cues exactly indicated upcoming effort levels. Effort cues were either red or green (blue in the figure) circles in Exp. 1 and upward or downward triangles in Exp. 2, and cue types were counterbalanced across participants. Left and right arrows probabilistically determined which types of effort cues were presented. The probabilities of arrows associated with effort cues, which indicated upcoming effort levels exactly, changed dynamically over time (Figure 1B). The set of either 80 or 20% was exchanged 5 times during 180 trials. Following arrow cue selection, an unambiguous effort cue thus resolved uncertainty about upcoming effort demands. There was no time limit for response. (B) An example of the presentation rates of low-demand problems (moving average of latest 5 trials) associated with the left arrow-cue (light grey) and the right arrow-cue (dark grey).

## Results

### Summary of experimental designs

We conducted two fMRI demand-selection experiments using mental arithmetic and spatial reasoning tasks ((Experiment (Exp.) 1: n=30 (males: females = 17:13), Exp. 2: n=28 (males: females = 12:16)) (Figure 1A). Both experiments had identical task structures but differed in cognitive demands. At the start of each trial, participants chose either a left or right arrow. After the arrow choice, one of two effort cues informing participants of upcoming effort demands was presented to participants, based on their choice. Choices transitioned probabilistically to either high or low effort cues (Figure 1A). These transition probabilities changed across 180 trials with five reversals (Figure 1B). Participants were instructed “the probability that each type of problem is more likely to appear is set independently for the left and right arrows. These probabilities change slowly over time. The same situation will continue for at least 10 trials”. After the effort cue, a problem was presented. Participants were instructed to “solve the problem as fast and accurately as possible and redo the task if you think the answer may be wrong”. Participants did not receive feedback on their response accuracy. There was no time limit either for arrow choices or answers to problems. We instructed that choices did not affect session duration.

For both experiments, we prepared two sets of problems that required different levels of cognitive demand: low- and high-demand problems. The rule of mental arithmetic (division) problems of Exp. 1 was to divide a 5-digit number by 7 and report whether the remainder was small (<= 3) or large (>= 4). The dividend in low-demand problems (e.g., 35426) consisted of “two consecutive two-digit numbers that were multiples of 7” followed by a single one-digit number from 1 to 6, whereas the dividend in high-demand problems (e.g., 48106) did not contain any numbers that were multiples of 7. The rule of spatial reasoning (mental cube-folding) problems of Exp. 2 was to judge whether a 3D cube with three colored faces and a simultaneously presented unfolded cube matched or not. The demand difference in spatial reasoning problems was whether the three faces shown on the 3D cube were adjacent on the unfolded cube (low-demand) or not (high-demand).

### Answers of post-experimental questionnaire

In Exp. 1, 97% of participants, and in Exp. 2, 100% answered “yes” when asked whether the instruction was consistent with their scanner experience. These results suggest that most participants thought that the probabilities of two problem types changed independently, though, in reality, the probability with which either arrow transitioned to each effort cue (80% or 20%) changed five times during 180 trials.

We also asked how much participants liked to solve either one of two problems. The mean score on a 7-point scale was 1.00 ± 0.18 (mean ± SEM) in Exp. 1 and 0.93 ± 0.22 in Exp. 2 (0 = easiest, 6 = most difficult). The individual differences of this avoidance scales did not correlate with individual differences of adjusted standardized residuals of a chi-square test which we use to index avoidance in either experiment (r = 0.18, p = 0.34 in Exp. 1 and r = 0.35, p = 0.07 in Exp. 2). These results suggest that most participants avoided high-demand problems but stated preferences and choice tendencies to avoid high-demand problems did not strongly relate.

### Response time of arrow choices and task performance

The response time for choosing an arrow cue (RT_choice_) was 1.19 ± 0.12 sec (mean ± SEM) in Exp. 1 and 1.06 ± 0.08 sec in Exp. 2. The mean accuracies for high- and low-demand problems were, respectively, 0.95 ± 0.014 (mean ± SEM) and 0.98 ± 0.005 in Exp. 1, and 0.92 ± 0.014 and 0.99 ± 0.002 in Exp. 2. In both experiments, the rate of correct answers to the low-demand problems was on average higher than the rate of correct answers to the high-demand problems (paired t-test, t (29) = -3.10, p = 4.3 ×10^-3^, Exp. 1, t (27) = - 5.79, p = 3.7 × 10^-6^, Exp. 2). The mean solution times for correctly answered high- and low-demand problems were, respectively, 14.72 ± 1.22 sec (mean ± SEM) and 1.91 ± 0.16 sec in Exp. 1, and 12.91 ± 0.84 sec and 4.11 ± 0.18 sec in Exp. 2. In both experiments, participants took longer for the high-demand problems (paired t(29) = 11.02, p = 7.0 × 10^-12^, in Exp. 1, t(27) = 11.94, p = 2.8 × 10^-12^, in Exp. 2). We set inclusion criteria (> 80% accuracy for all main experiment trials of both high- and low-demand problems) to ensure engagement. By this criterion, we excluded one and two participants in Exp. 1 and Exp. 2 from further analyses, respectively.

### Avoidance Behavior

We assumed that participants who tried to avoid high-demand problems would stay on the same side (left or right) after solving a low-demand problem in the previous trial and would switch to the opposite side after solving a high-demand problem. We tested this with a chi-square test (2 × 2 factors: problem type [high- or low-demand] in the previous trial × choice [same side or opposite side] in the current trial), as same as our previous study^11^. We found that 27 (93.1 %, out of 29) and 24 (92.3%, out of 26) participants in Exp. 1 and Exp. 2, respectively, significantly tended to avoid high-demand problems (p < 0.01), indicating that most are demand avoiders. These results indicate that, reflecting on experiences one trial back, most participants learned to avoid high-demand problems in the uncertain environment where the upcoming effort level changed over time. Most participants avoided high-demand problems (Chi-square test, p < 0.01)^11^.

### Adaptive learning to avoid mental effort with reflection on the experiences of two preceding trials in demand avoiders

We tested the effects of experiences two trials back for demand avoiders who significantly avoided high-demand problems (chi-square test). We categorized choices into four bins: problem type (high- or low-demand) in the (k–2)-th trial × problem type in the (k–1)-th trial for trials in which the (k–1)-th choice was the same or opposite of the choice on the (k–2)-th trial. We found that the proportions of the k-th choice that were the same as (k– 2)-th choice depended on both choice and experience history (Figure 2ABC). There were some significant differences in the means: participants chose the same options with the (k–2)-th choice more frequently when the experienced demand of two trials-back was low versus high, with small to large effect sizes in Exp. 1 and 2, respectively (Figure 2Aa: p = 1.7 × 10^-8^, d = -1.79, t (26) = -8.01; Figure 1Ac: p = 2.6 × 10^-2^, d = -0.74, t (22) = -2.40; Figure 2Ad: p = 4.7 × 10^-2^, d = -0.56, t (22) = -2.10) and in Exp. 2 (Figure 2Ba: p = 2.6 × 10^-8^, d = -1.49, t (23) = -8.24; Figure 2Bc: p = 4.1 × 10^-4^, d = -1.08, t (22) = -4.15). Most participants significantly avoided high-demand problems, adapting based on experiences from two preceding trials in a changing environment. These results all replicate our previous findings^11^ that humans avoid mental effort based on a demand history.

### Demand avoiders reflected multiple past experiences to their current choices

Using Generalized Linear Mixed Model (GLMM) analyses, we tested whether one, two, and more than three past experiences affect the current choices of demand avoiders. The model included fixed effects of the choice and outcome from (k–1)-th to (k–4)-th trials, as well as the (k–1)-th choice. We assumed random effects to properly allocate variance to between- and within-subjects predictors. The random effects were the individual variations in intercepts, in the choice and outcome of both (k–1)-th and (k–2)-th trials, and in (k–1)-th choice. Choices were significantly influenced by the choice and outcome of (k–1)-th, (k–2)-th, and (k–3)-th trials, and the choice at (k–1)-th trial significantly affected the current choices in both Exp. 1 and 2 (Figure 2D: logit coefficients 1.55, 0.72, 0.20, and 1.50, p < 0.05, in Exp. 1; Figure 2E: logit coefficients 1.73, 0.85, 0.30, and 1.58, p < 0.05, in Exp. 2). The choice and outcome at the (k–4)-th trial was also significant in Exp. 1 (Figure 2D: the logit coefficient was 0.23, p < 0.05), but not in Exp. 2 (p = 0.156). These results demonstrated that choices reflect multiple past experiences. More recent experiences had greater influence, suggesting a RL-like process.

**Figure 2.**
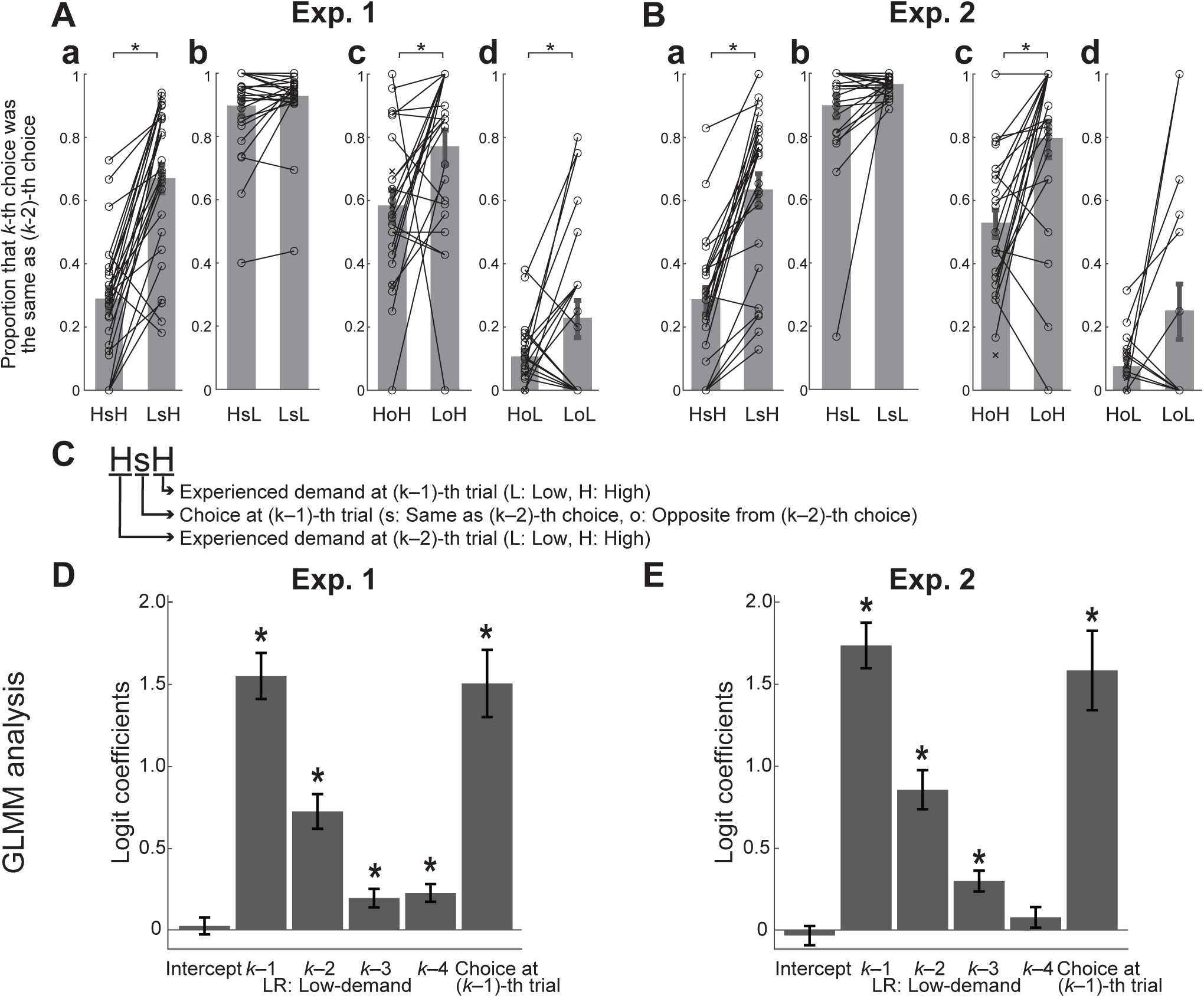
Demand avoiders made a choice, learning based on multiple past experiences. Behavioral analyses for the choices of demand avoiders in Exp. 1 (n = 27) and Exp. 2 (n = 24). (AB) The influences of the two preceding trials on current choices. Each pair of bars indicates the across-participants mean rates that the current (k-th) choice was the same as the choice at 2 trials before ((k–2)-th) when the experienced demand at the (k– 2)-th trial was high (left bar) or low (right bar), for each case sorted by the choice at the (k–1)-th trial (same as (a, b) or opposite from (c, d) the (k–2)-th choice) and the experienced demand at the (k–1)-th trial (high (a, c) or low (b, d)). Each bar has a label like HsH, following the rule explained in (C). The average was taken across the participants who were judged as demand avoiders as a result of Chi-square test (see Result section). The error bars indicate the mean ± standard error of the mean (SEM). The paired data of individual participants were shown as dots and were connected by lines. Unpaired data were shown as crosses without lines. The paired data were compared by paired t-test, and asterisks marked significant differences. (C) Bar label explanation. The HsH is an example of bar labels on Figure 2A and 2B. (D,E) The influences of multiple past experiences on the current choices, analyzed using Generalized Linear Mixed Model (GLMM). Bars indicate the fixed effects and the SEMs. The fixed effects were an intercept, influences of the choice and outcome from both the (k–1)-th to the (k–4)-th trials and the choice at (k–1)-th trial on the choice at k-th trial. Random effects were the individual variations in intercepts, in the choice and outcome of both (k–1)-th and (k–2)- th trials, and in the choice at (k–1)-th trial. Asterisks mark the significant t-test at p < .05.

### Model-based analysis of choice data

We performed model fitting and comparison to determine which models best explain choice behavior in effort avoidance. Eleven models were tested, incorporating cost functions based on solution times, inaccuracy rates, rest time, and binarized demand levels (see Methods and Mitsuto et al., 2018 for full details)^11^. Nearly the same models were used as in our prior study, though we cut off the five worst-performing models. Bayesian Information Criterion (BIC) was used for model comparison. RL models assumed participants learned each factor in a simple RL update algorithm based on the prediction errors. We assumed Solution-Times (ST; continuous), Incorrect-Correct (IC; binary), and High-Low demand levels (HL; binary) as *ActualCost* in RL models. We also tested ST-IC and HL-IC combination models. Another model assumed rial-by-trial rest-time (Rest) as a reinforcer and tested the combination between RL-HL and Rest. We also tested one and two-parameter probabilistic models (Win-Stay-Lose-Shift: pWSLS and Probabilistic-Selection: PS), assuming participants chose strategically based on HL or IC (HL and IC: binary). In the pWSLS-HL, a participant stays with their choice after a low-demand problem and switches after solving high-demand problems with the probability *p*. In the PS-HL model, the probabilities were *a* and *b* (the repeat rate after high and low demand, respectively).

Among RL models, the RL-HL model (binarized demand) had the best BIC scores for most participants in Exp. 1, and all the participants in Exp. 2 (25/27 (92.6%) in Exp. 1; 24/24 (100.0%) in Exp. 2; Figure 3Aa and Figure 3Ba, Table S1). The result that the RL-HL model outperformed the other RL models indicates that the cost of mental effort is learned as a binary variable, rather than variables reflecting the solution time or errors. The RL-HL model was also better than the two rest-time RL models (27/27 (100.0%) in Exp. 1; 21/24 (87.5%) in Exp. 2; Figure 3Ab and Figure 3Bb). Among probabilistic models, the PS model with binarized effort demands was superior (25/27 (92.6%) in Exp. 1; 23/24 (95.8%) in Exp. 2; Figure 3Ac and Figure 3Bc). Direct BIC comparisons showed that the PS model outperformed RL-HL (17/27 (63.0%) in Exp. 1; 14/24 (58.3%) in Exp. 2). Thus, both PS and RL models captured effort avoidance, but PS explained choices better for more participants.

Decisions in the PS model rely on one preceding trial, while RL models incorporate multiple past trials (see Methods). According to BIC values, the PS model explained choices better than the RL model in both our previous and current studies. On the other hand, the RL model captured the sequential effect of past experienced demands that the PS model was incapable of capturing. Previous simulations showed that RL, but not PS, captured the effect of two-preceding trials (Figure 8 of Mitsuto et al.^11^). Thus, we conducted model-based fMRI analysis using the RL-HL model. Figure 3C presents a sample RL-HL model fitting. Figure 3D shows estimated CPE values of the same sample data. Model-based fMRI analyses were conducted using CPE values from the RL-HL model.

**Figure 3.**
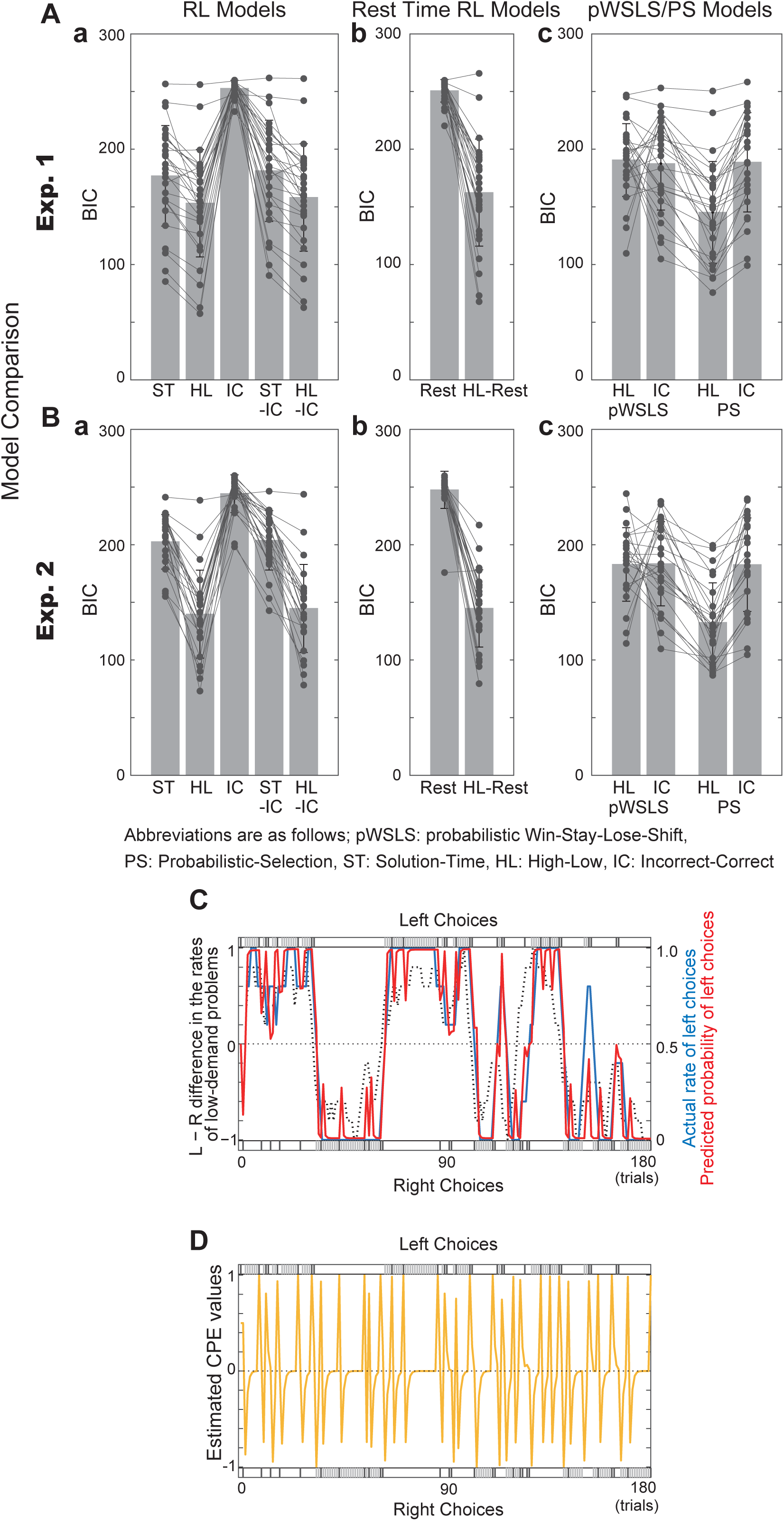
Model comparisons, examples of model fitting and the estimated cost prediction error values. (A,B) Model Comparison. The bars indicate the mean ± SEM of the Bayesian Information Criterion (BIC) scores. Results for 5 models of reinforcement learning (RL) based on ActualCost (Solution-Time (ST), High-Low (HL), Incorrect-Correct (IC), ST-IC, and HL-IC). Individual demand avoiders are represented by the black dots linked with lines. (Ab, Bb) Results for the rest-time RL models which regarded the rest time in the scanner as reward for participants. (Ac, Bc) Results for probabilistic Win-Stay-Lose-Shift (pWSLS) models and Probabilistic-Selection (PS) models based on HL and IC. (B) Illustration of choices and choice probability predicted by the reinforcement learning model based on High-Low demands (RL-HL model). Short vertical bars at the top and bottom indicate participant’s left and right choices, respectively, with dark- or light-gray indicating that high- or low-demand problems were experienced, respectively. The black dashed line in the middle indicates the Left – Right difference in the presentation rates (moving average of latest 5 trials) of low-demand problems plotted against the left scale. The blue and red solid lines indicate the participant’s actual left-choice rate (moving average of latest 5 trials) and the left-choice probability predicted by the RL-HL model plotted against the blue and red scales on the right, respectively. (C) Illustration of estimated values of cost prediction error of the same participant with Figure 3B.

### Lack of CPE encoding at effort cues

We initially hypothesized that effort costs are learned by TD learning and expected costs are updated at the time of effort cues. We thus predicted that CPE signals correlate positively with dmFC/dACC, and negatively with striatal BOLD signal, irrespective of demand type, at the time of effort cues (Figure 4Aa). To test this hypothesis, we examined the activity of the whole brain to search for regions where BOLD signal at effort cue correlated positively or negatively with the CPE signals in a conjunction analysis between Exp. 1 and Exp. 2. The conjunction analysis involved a strict mask comprising common voxels from Exp. 1 and 2, with the threshold of cluster-level FWE corrected p < 0.05, and voxel-level uncorrected p < 0.001 from GLM 1 (Figure 4Ba). At the time of effort cues, activity in neither the dmFC/dACC nor the striatum were correlated with CPE signals significantly with the threshold of cluster-level FWE corrected *p* < 0.05, and voxel-level, uncorrected *p* < 0.001 (Figure 5A, Figure S1ABGH, Table S2; the results of conjunction analyses were reported with this threshold, unless otherwise stated). Given the conservative nature of this test, we used a relaxed mask. To confirm the null results, we analyzed the conjunction using a relaxed mask that consisted of common voxels between the results of Exp. 1 and 2 with the threshold of voxel-level uncorrected p < 0.01 using GLM 1. Under relaxed criteria, activity in the vmPFC/vACC was positively correlated with the CPE signals at the time of effort cues (voxel-level uncorrected p < 0.001, Table S2D), but there was no super-threshold correlation with activity in the dmFC/dACC. These results thus fail to support the first hypothesis (Figure 4Aa).

**Figure 4.**
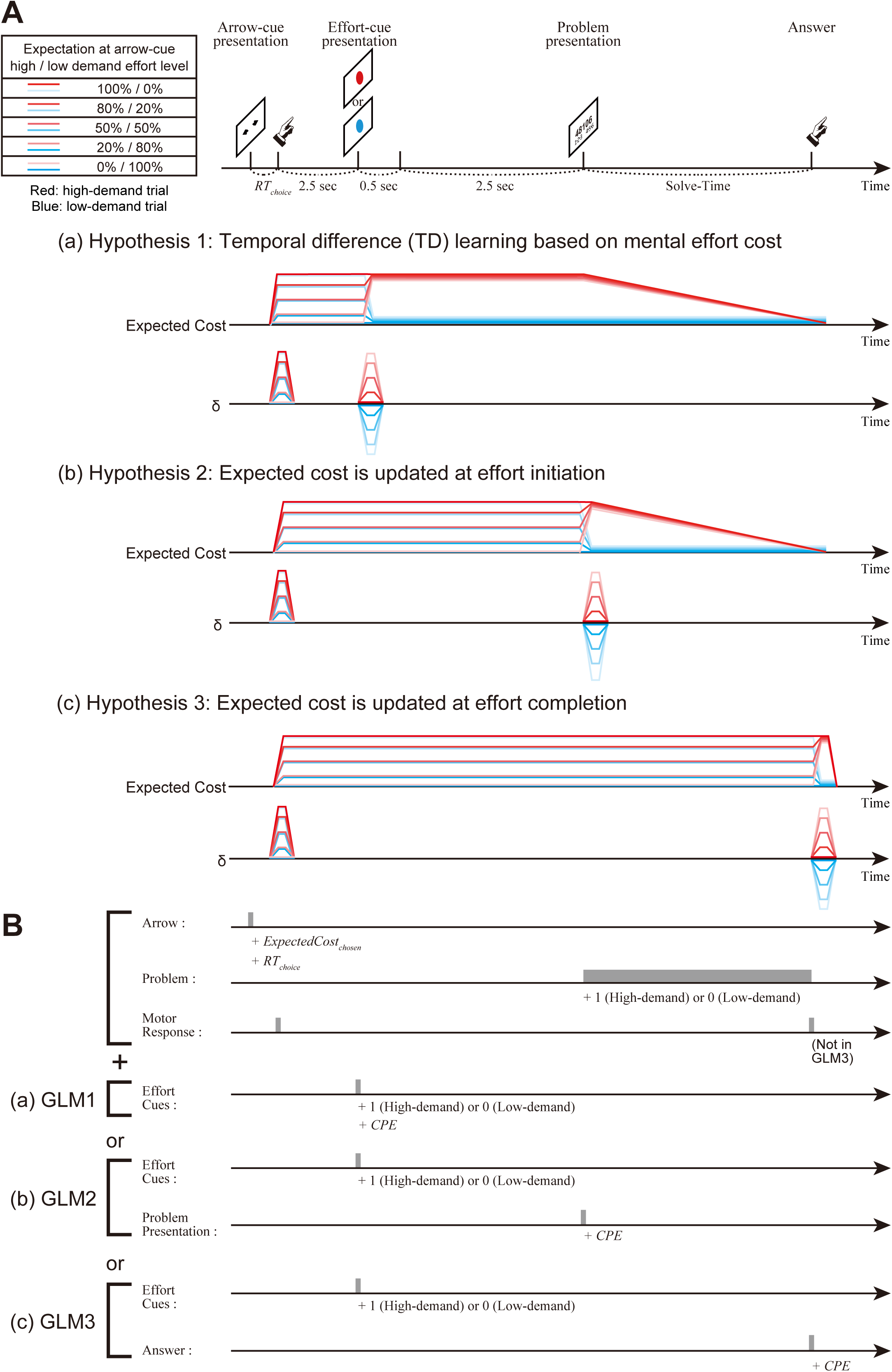
Hypotheses and General Linear Models for fMRI Analyses. (Aa) Hypothesis 1 is that effort cost learning follows temporal-difference learning such that expected costs are updated at the time of the effort cues. If participants predict the upcoming effort demand is high-demand at 80% and they receive the information of low demand at the effort cue presentation, the prediction error (δ) of -80% generates at the effort cue timing. Expected cost and the cost prediction error (CPE) as δ during a trial are shown schematically. Lines are colored for high-demand trials with the gradation from red to pale pink and for low-demand with the gradation from pale blue to blue, representing expected effort levels at the arrow-cue from high- to low-demand effort levels. To simplify, we used a monotonic linear decrease function as mental effort cost function during problem solution. (Ab) Hypothesis 2 is that expected costs are updated at effort initiation. (Ac) Hypothesis 3 is that expected costs are updated at effort completion. (B) There were five events in a trial: presentation of arrow cues, arrow cue choice, presentation of an effort cue, problem presentation, and problem completion. We explored the neural correlates of the CPE estimated in the reinforcement learning model based on effort levels by using three GLMs (GLM1-3), according to three different hypotheses regarding the time of CPE generation. All GLMs included regressors at the presentation of arrow cues with non-orthogonalized parametric modulations by ExpectedCostchosen and RTchoice, regressors at presentation of effort cues with non-orthogonalized parametric modulations by upcoming demand-level (1 and 0 for high- and low-demand problems, respectively), regressors at arrow cue choice, regressors problem presentation having the duration of solution-time with non-orthogonalized parametric modulations by demand-level (1 and 0 for high- and low-demand problems, respectively). Respectively, the GLMs also contained regressors for motor response at both arrow choice and answer (GLM1 and 2) or arrow choice (GLM3), regressors for head movements (not illustrated here), and CPE-modulated regressors at effort cue ((a) GLM1), at problem onset ((b) GLM2), or at answer time ((c) GLM3).

**Figure 5.**
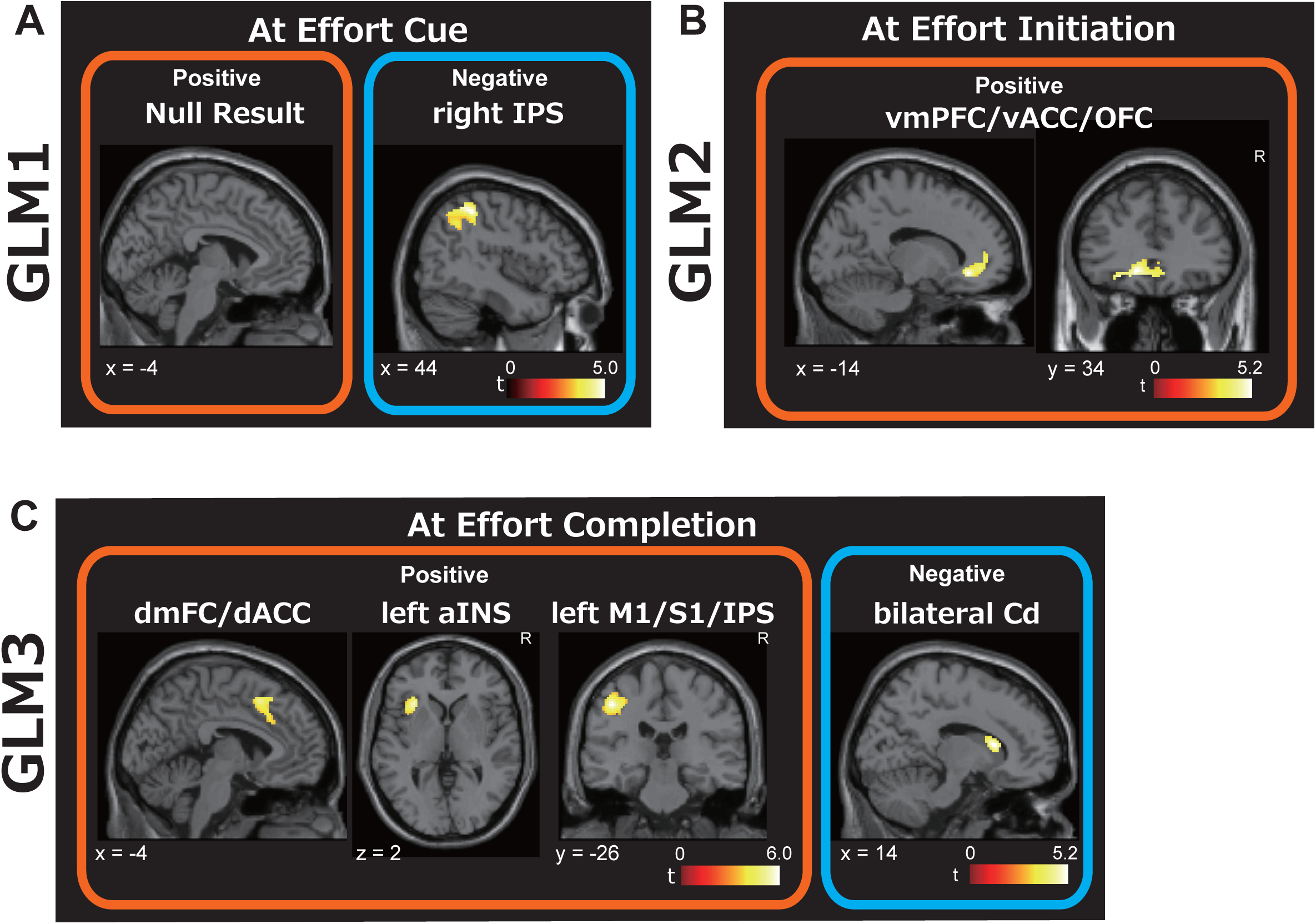
Neural correlates of the cost prediction error at the time of effort cue, at effort initiation, and at effort completion. Conjunction analyses with a binary mask consisting of common voxels between the positive and/or negative correlations of the cost prediction error (CPE) at the time of effort cue (A: GLM1), problem presentation (B: GLM2) or answer (C: GLM3) in Exp. 1 and Exp. 2 at cluster-level FWE corrected p < 0.05 and voxel-level uncorrected p < 0.001. (A) No clusters were detected as a positive correlation. The right inferior parietal sulcus (IPS) was detected as a negative correlation. (B) The vmPFC/vACC/orbitofrontal cortex (OFC) was detected as positive correlation. (C) Three clusters were detected in the dorsomedial frontal cortex/ dorsal anterior cingulate cortex (dmFC/dACC), left anterior insula (aINS), left primary motor cortex (M1)/ primary somatosensory cortex (S1)/IPS as a positive correlation. The bilateral caudate nucleus was detected as a negative correlation.

In contrast to these null results, activity in the right inferior parietal sulcus (IPS) correlates negatively with CPE signals at the time of effort cues, using the strict mask (Figure 5A, Figure S1GH, Table S2). This finding is discussed later.

One possibility for the null results is that participants ignored effort cues. However, conjunction analysis showed significant task preparation responses in dmFC/dACC, right dlPFC, and right Angular Gyrus (AG) activity, which positively correlated with problem demand (Figure 6). This result indicates that most demand avoiders encoded information about upcoming effort demands at the time of effort cues, predicting the upcoming effort, and prepared for the task accordingly. Therefore, these null results imply that the brain does not encode CPE signals to update expected costs at the moment when full information becomes available and, as such, mental effort cost learning does not follow traditional TD learning.

**Figure 6.**
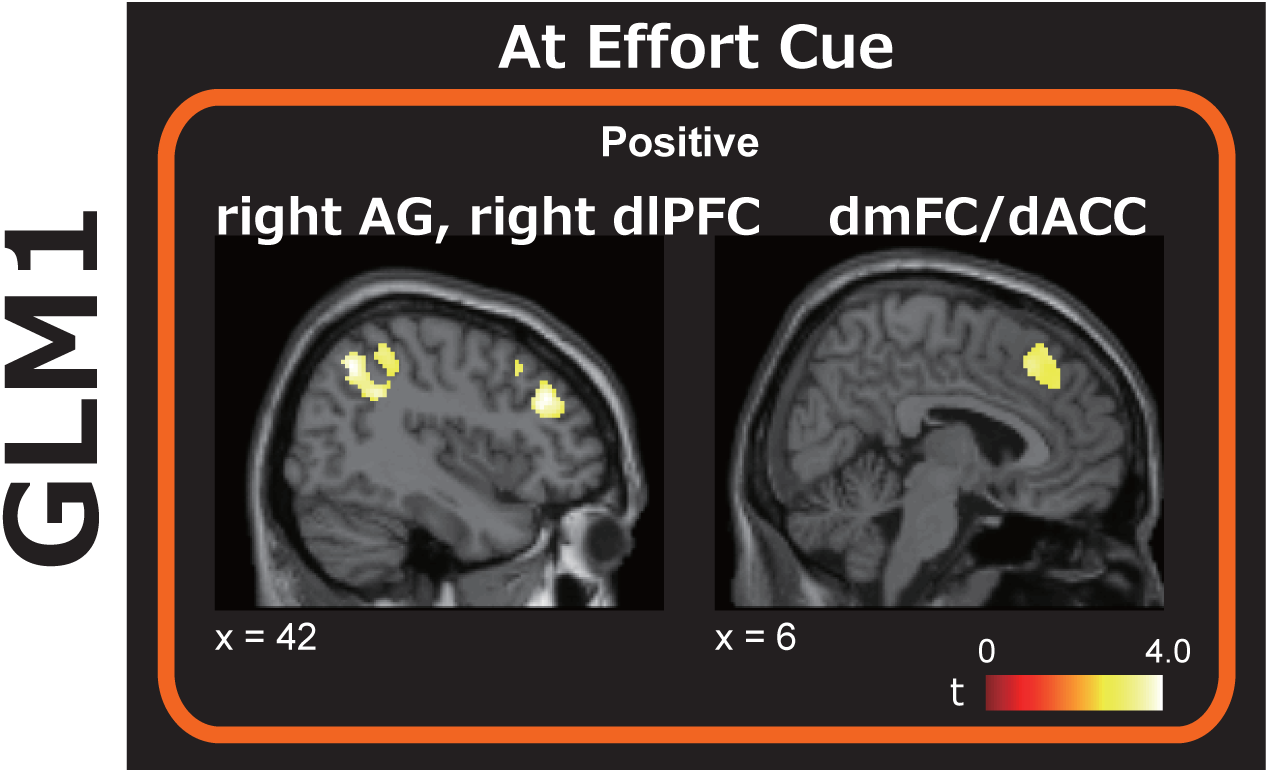
Neural correlates of the task preparation at presentation of an effort cue. Conjunction analyses with a binary mask consisting of common voxels between the positive correlations of the task preparation at the time of effort cue (GLM1 was used) in Exp. 1 and Exp. 2 at cluster-level FWE corrected p < 0.05 and voxel-level uncorrected p < 0.001. The right dorsolateral prefrontal cortex (dlPFC), right angular gyrus (AG), dmFC/dACC were detected as a positive correlation.

### CPE encoding at effort initiation

We next hypothesized that expected cost updates require effort initiation (Figure 4Ab). A conjunction analysis with the strict mask by using GLM 2 (Figure 4Bb) revealed that, at the time of problem presentation when effort is initiated, the activity in the ventromedial prefrontal cortex (vmPFC)/ ventral anterior cingulate cortex (vACC)/orbitofrontal cortex (OFC) correlates positively with CPE signals (Figure 5B, Figure S1CDIJ, Table S3). Although we detected a positive correlation between the CPE and the activity of the large cluster including the dmFC/dACC in Exp. 1, the CPE was neither positively correlated with the activity in the dmFC/dACC in Exp. 2 nor negatively correlated with the activity in the striatum in either experiment. These results fail to support the second hypothesis (Figure 4Ab). However, they suggest that the vmPFC/vACC/OFC encode CPEs, at the time of effort initiation, with the information about upcoming effort levels.

### CPE encoding at effort completion

We next tested the hypothesis that expected costs are updated at effort completion (Figure 4Ac). A conjunction analysis with the strict mask (GLM 3; Figure 4Bc), revealed that activity in the dmFC/dACC, the left anterior insula (aINS), and the left primary motor cortex (M1)/ primary somatosensory cortex (S1)/IPS correlates positively, and activity in the bilateral caudate nucleus (Cd) correlates negatively with CPEs at the time of effort completion (Figure 5C, Figure S1EFKL, Table S4). Therefore, these results support the third hypothesis and support the hypothesis that expected costs are updated at the time of effort completion (Figure 5C).

### Residual BOLD signals at effort completion predicted sequential choices

If CPE encoding reflects an update to choice policy, then BOLD signal at the time of CPE encoding should predict choice on the next trial. We tested the prediction that averaged residual BOLD signal extracted from each cluster ROI that encoded CPEs anticipates subsequent stay-switch choices, using mixed effects logistic regressions. For our analysis, we first fit a new GLM, identical to the model we fit previously, but without a CPE regressor (Figure 4B – without +*CPE*), and extracted the residual BOLD signal from all ROIs defined by our prior conjunction analysis to correlate with CPEs at various time points. Finally, we regressed binary stay-switch choices onto extracted residuals for each ROI. In this analysis, a positive (negative) regression weight indicates that a participant is more (less) likely to switch their choice on the next trial, reflecting an update in choice policy. We find that residual BOLD signals in the Cd at effort completion predicted sequential choices significantly in both Exp. 1 and 2 (Figure 7A). The residual BOLD in the dmFC/dACC, left aINS, and left M1/S1/IPS also predicted the upcoming choices in Exp. 2 (Figure 7AB). Neither BOLD signal residuals in the right IPS at effort cues nor the vmPFC/vACC/OFC at effort initiation predicted upcoming choices (Figure 7A, Figure S2AB), though all clusters’ residual BOLD signals were correlated with CPEs (Supplemental Result 1). These results imply that the neural signals at effort completion reflect an update to choice policy while CPE-correlating activity either at effort cues or at effort initiation does not.

**Figure 7.**
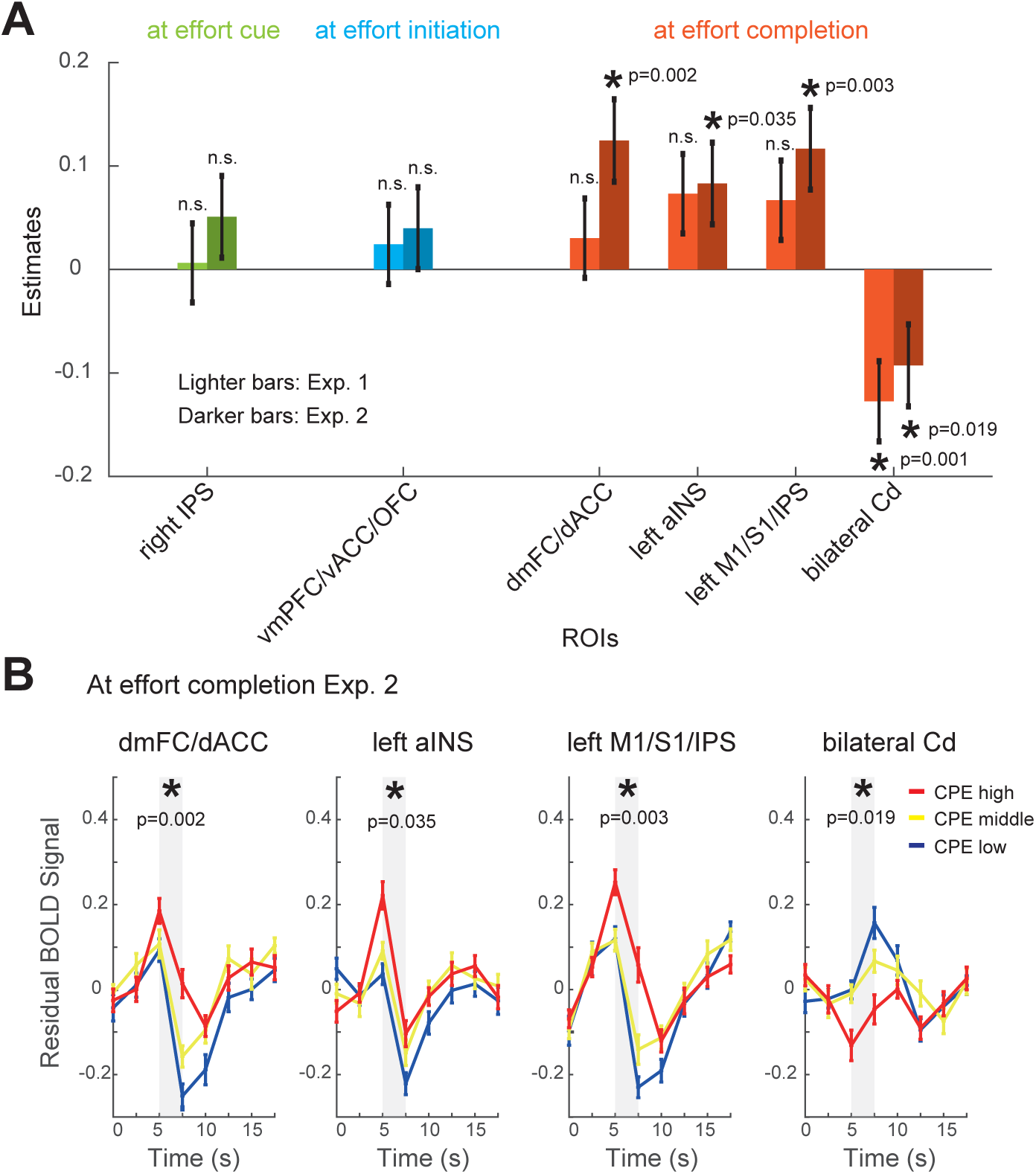
Cluster ROI-based residual BOLD signal analyses: Prediction of sequential choices. (A) The influence of residual BOLD signals from each cluster region of interest (ROI) for each time point (effort cue: green bars, effort initiation: blue bars, and effort completion: red bars) on subsequent choices, analyzed using a Generalized Linear Mixed Model (GLMM). Fixed effects included the residual BOLD signal at the k-th trial to predict subsequent choice behavior (switch or stay). Random effects accounted for individual variation in the intercepts. Asterisks mark significant t-tests at p ≤ 0.05. Each p-value was shown. Lighter bars indicate results from Experiment 1, while darker bars indicate results from Experiment 2. Positive estimates suggest a higher residual BOLD signal is associated with an increased likelihood of switching choices in subsequent trials, while negative estimates indicate an increased likelihood of staying. (B) Time-course analyses of residual BOLD signals at effort completion (Experiment 2) for selected ROIs: the dmFC/dACC, left aINS, left M1/S1/IPS, and bilateral Cd. Curves represent residual BOLD signals averaged by CPE tertiles: CPE Low (−1 ≤ CPE < 33rd percentile, blue line), CPE Middle (33rd ≤ CPE ≤ 67th percentile, yellow line), and CPE High (67th percentile < CPE ≤ 1, red line). Before taking group averages, the residual BOLD signals for each participant were normalized in two steps: first, within each block using z-scores, and second, across all blocks for each subject using z-scores. Shaded regions indicate time points (5s and 7.5s) where the averaged values were used for analysis. Asterisks mark significant t-tests at p ≤ 0.05 in the GLMM.

## Discussion

We first hypothesized that mental effort-cost learning follows TD learning and that cued information about upcoming effort is sufficient to trigger an update of expected effort associated with a given choice. We therefore predicted that, at the time of effort cues, CPEs correlate positively with activity in the dmFC/dACC and negatively with activity in the striatum. However, activity in neither the dmFC/dACC nor striatum correlate with CPEs at the time of effort cues, which was revealed by a conjunction analysis across Exp. 1 and 2. This result failed to support the first hypothesis. Therefore, we hypothesized that expected costs are updated either at effort initiation or completion. At effort initiation, the activity of the vmPFC/vACC/OFC correlates positively with CPEs. At effort completion the CPE correlates positively with the activity in the dmFC/dACC and other regions, and correlates negatively with activity in the bilateral Cd. Additionally, we tested whether such correlates of CPE reflect an update of effort expectations, and thus choice policy. Thus, we tested the prediction that BOLD signal in regions that encode CPEs also predict choice on the next trial. We found that residual BOLD signals at effort completion predicted the sequential choices in both Exp. 1 and 2. These results demonstrate that mental effort cost learning requires effort exertion and imply that adaptive learning of mental effort costs does not follow traditional TD learning.

### Consideration of results and limitations

#### Multiple Past Demanding Experiences Affect Current Choices

Behavioral data showed that humans tend to avoid mental effort, reflecting the experiences of multiple past events in the task design of the current study with the addition of effort cues, replicating our previous study^11^. The choice data of demand avoiders were approximated with a RL model based on demand levels, implying that humans track mental effort cost information. Questionnaire results suggested that most participants prefer low-demand problems but self-reported demand preferences have a weak relationship with behavioral choice tendencies.

#### Lack of Significant Results at Effort Cues

Activity in neither the dmFC/dACC nor striatum correlated with CPEs at the time of effort cues. This lack of a CPE correlation with the activity in the dmFC/dACC when effort information is first available contradicts the prediction from the TD hypothesis (Figure 4A and 5A). Furthermore, residual BOLD in the right IPS at effort cue did not predict the sequential choices, and the residual BOLD signals in right IPS did not show a consistent activity pattern with respect to CPEs between two experiments (Figure S2A). We speculate that the negative correlation between right IPS activity and CPE reflects a network reset signal in the frontoparietal network at effort cue timing^32^. Consistent with the network reset hypothesis, increased IPS activity in response to a negative CPE (indicating an easier-than-expected task) may reflect disengagement from a high-attentional state. Collectively, our results are inconsistent with the hypothesis of traditional TD learning whereby mental effort cost expectations are updated when effort information first becomes available. We acknowledge the possibility that our methods are insufficiently sensitive to detect CPE encoding in the dmFC/dACC and the striatum at informative effort cues. Indeed, the absence of evidence is not the evidence of absence. However, our methods were sufficiently sensitive to detect CPEs at effort completion, suggesting at the very least that CPE encoding is more robust at effort completion.

#### vmPFC/vACC/OFC Activity at Effort Initiation

Activity patterns around the rostromedial prefrontal cortex were consistent across two experiments in our previous study that did not use effort cues^11^ and four experiments in total. However, the positive correlation between CPEs and activity in vmPFC/vACC/OFC at effort initiation contradicts the second hypothesis that expected costs are updated at the time of effort initiation (Figure 4Ab and 5B). These results stand in contrast with the general perspective in the field of value-based decision-making that the vmPFC positively encodes predicted reward value, or reward PEs, discounted by effort costs^31,33–35^. A positive correlation with costs thus argues against the prediction that the vmPFC activity at effort initiation reflects value encoding or value updating. Furthermore, residual BOLD signals of the vmPFC/vACC/OFC at effort initiation do not predict the subsequent choices in either experiment, implying that although BOLD signal correlates with CPEs, this correlation does not reflect an update of expected costs that subsequently guides value-based decision-making.

Activity in the vmPFC/vACC/OFC also correlated with CPEs at effort initiation, which we speculate reflects a shift to a task engaged state. More generally, the OFC is hypothesized to track current state for elaborating progress through a model of the external world, cognitive map^36,37,38,39^. The OFC may track current states and update a cognitive map such as state-space map and task set during effort initiation, supporting transitions into task-engaged states and credit assignment for actions directed toward task goals. Perhaps the mental action, triggered by effort initiation, recruits the OFC to update a world model, reflecting a transition into task-engaged states.

#### Sequential Choice Predicted by Residual BOLD Signal at Effort Completion

When participants complete a task, the associated CPE correlates positively with the activity in the dmFC/dACC, left aINS, and left M1/S1/IPS and negatively with activity in the bilateral Cd, irrespective of demand type (Figure 4Ac and 5C). Furthermore, residual BOLD signals predicted the upcoming choices in the Cd, in both experiments, as well as the residual BOLD signals at the dmFC/dACC, left aINS, and left M1/S1/IPS clusters in Experiment 2 (Figure 7AB, Figure S2C). These results demonstrate that the brains encode the CPE signals at effort completion, not at other timepoints, and moreover that this CPE encoding at effort completion reflects an update of expected values and choice policy.

Potential confounds, such as retrospective confidence PE or satisfaction PE, were considered. However, confidence-related activations are typically observed in regions like the frontopolar cortex or ventrolateral PFC, not in the Cd^40–42^. Similarly, satisfaction as a confound is unlikely since it serves as a positive reinforcer. Because participants avoided higher demands, they did not behave as though they found greater satisfaction when completing higher demands.

#### Mental Effort Learning Algorithm

We speculate that expected cost updates are delayed until task completion because people can adjust effort on-the-fly to handle unexpected difficulties or unexpected efficacies. With practice and insight, some tasks may become easier, so the brain may wait to evaluate mental effort retrospectively, after completion, to access full information about the effort that was just exerted. Waiting until effort completion also avoids multi-tasking between the task itself and the evaluation of effort costs.

Our data imply that mental effort costs are learned in cortico-basal ganglia loops encompassing the dmFC/dACC and Cd, being consistent with known circuit anatomy^27^. This hypothesis is supported by evidence of positive encoding of effort costs and effort preparation in the dmFC/dACC and the negative correlation of the CPE with activity in the Cd, detected at effort completion.

Traditional TD learning theory failed to explain our results. Recently, the retrospective causal learning algorithm was proposed to account for midbrain dopamine signaling, during reward learning, better than classic TD learning^43^. While our results indicate that mental effort learning entails retrospective inference, the retrospective causal learning algorithm does not explain why CPEs are encoded at effort completion. As with TD learning, retrospective causal learning also predicts a CPE when information is first available. Therefore, we propose that the algorithm governing mental effort cost learning involves retrospectively Bayesian or non-TD RL^1,44^, updating expected effort costs as a function of the difference between prior expectations, and experienced effort costs, reported as a CPE, when exertion is completed.

### Future Directions and Perspectives

Refining the effort cost learning algorithm is essential. Future studies should simulate and test multiple candidates learning algorithms, based on the observation that updates require direct experience, to develop new experimental paradigms and determine which algorithm best explains learning.

## Conclusion

Our data indicate that mental effort cost learning is retrospective and that the learning algorithm of mental effort costs is not classical TD learning. We find that humans update expected costs by CPE when they finish exerting mental effort. Furthermore, we find that the residual BOLD signals at effort completion predicted subsequent choices, indicating that the CPE encoding reflects an update of choice policies at the time of effort completion. Thus, effort exertion is required for humans to learn the costs of effortful tasks. These findings refine our understanding of the mechanisms underlying decision-making and learning based on effort with implications for adaptive learning of mental effort in health and disease.

**Supplemental Result 1. Residual BOLD signal of each cluster region of interest (ROIs) explained CPEs.** To confirm the trial-wise relationship between each averaged residual BOLD signal of each cluster ROIs at each timing and CPE signals, a Linear Mixed Model (LMM) analysis was conducted. The cluster ROIs at each time point were as follows: the right IPS at effort cue, the vmPFC/vACC/OFC at effort initiation, and the dmFC/dACC, left aINS, left M1/S1/IPS, and bilateral Cd at effort completion. Residual BOLD signals within each ROI significantly explained CPE signals across trials.

## Author Contributions

1. Conceptualization AM
2. Data curation AM
3. Formal analysis AM
4. Funding acquisition SY, AW, and AM
5. Investigation AM, TK, and KO
6. Methodology AM, AW
7. Project administration AM, SY, and RH
8. Resources AM
9. Software AM
10. Supervision AM, SY, RH, and AW
11. Validation AM, AW, KM, TY, and KO
12. Visualization AM
13. Writing – original draft AM
14. Writing – review & editing AM, RA, KM, TK, RH, and AW

## Lead contact

Further information and requests for resources should be directed to and will be fulfilled by the lead contact, Asako Mitsuto.

## Acknowledgements

This work was supported by Grant-in-Aid of JSPS for Research Activity Start-up No. 18H06090 [to AM], for JSPS Fellows No. 19J00964 [to AM], for Early-Career Scientists No. 20K16475 and No. 23K14680 [to AM], Challenging Research (Exploratory) No. 19K21809 [to AM], and for Scientific Research on Innovative Areas No. 15H05876 [to AM], the Impulsing Paradigm Change through disruptive Technologies (ImPACT) program in Japan [to SY], and Grant No. R00 MH125021 of NIMH [to AW]. We are grateful to K Sakai, W Zajkowski (NIH), DL Aguilar (CBS), RP Badman (Harvard Univ), K Miyamoto (CBS), M Cai (Univ Miami), A Funamizu (Univ Tokyo), T Toyoizumi (CBS), Y Morishima (Bern Univ), K Tamura (Univ Cambridge), H Tomoki (ATR), A Nagai (Shimane Univ), T Shimizu (Tottori Univ) and the members of the Neurology Division of Tottori University, Akaishi Lab of RIKEN CBS, Doya Lab of OIST, and Rushworth Lab of the University of Oxford for the advice, support, discussion, or feedback to this study.

## Methods

### Participants

We recruited 30 participants (13 females; mean age, 23.0 ± 2.2) in Experiment (Exp.) 1 and 28 participants (16 females; mean age, 22.6 ± 1.6) in Exp. 2. Among these participants, 11 participants took part in both Exp. 1 and Exp. 2. We paid all participants 5,000 yen by bank transfer after their participation. No participants were taking any medicine or had a prior history of neuropsychiatric disorders. All participants were students of Shimane University, age 20-30 years, right-handed, native Japanese speakers, and received their compulsory education in Japan. Informed written consent was obtained from all participants before the experiment. The present study was approved by the medical ethics committee of Shimane University.

### Behavioral tasks

We conducted two experiments, Exp. 1 and Exp. 2 (Figure 1). Except for changes I∼VII as we described below, these experiments were almost similar behavioral protocols with our previous work ^11^. Consequently, we summarized the methods and listed changes I∼VII from our preceding procedure (please see Mitsuto *et al.*, 2018 for full details). The most important change was to add the effort cues that indicated the upcoming effort levels (i.e., high- or low-demand problems) between choices and problem presentations (please see the change I). There were two reasons why we added effort cues. First, if the first hypothesis was true and mental effort cost are learned via TD learning, this change should have allowed us to detect the CPE signals at the time of effort cues. This is because the PE signals generate to the earliest predictor, backpropagating from later predictors ^1^. In our previous design, the timing of the CPE generation was not locked by experimental manipulations and we did not know when the CPE signals were generated. Second, the addition of effort cues let us detect the signals of expected cost for the chosen option, reducing the confounding effect of task preparation at the time of choice-cues. At the time of choice-cues, the task preparation effects confound expected cost signals (expected effort demands) in both our previous and current designs. The effort cue reduced the necessity of the task preparation at the time of choice-cues in the current design because participants could get certain information of upcoming task type (i.e., high- or low-demand problems) at the time of effort cues, which was 2.5 seconds after the choice. We will report the results of the signals of expected cost of both Exp. 1 and Exp. 2 in another article.

We provided a summary of the experimental paradigms. The two experiments that we conducted had similar structures except for the following two points. The first point is that we used different types of problems in the two experiments. More specifically, we used mental arithmetic problems to divide a 5-digit number by 7 and report whether the remainder was small (<= 3) or large (>= 4) in Exp. 1, and spatial reasoning (mental cube-folding) problems to judge whether a 3D cube with three visible colored faces and a concurrently presented unfolded cube matched or not in Exp. 2. The second point is that we used different features of effort cues that indicated upcoming effort levels; circle cues with red and green colors in Exp. 1 and upward and downward triangle cues in Exp. 2.

There were problems with two levels of cognitive demand, i.e., high-demand and low-demand problems in both experiments. In Exp. 1, the dividend for the low-demand problem (e.g., 35426) was two consecutive two-digit numbers that were multiples of 7 followed by one single-digit number from 1 to 6, whereas the dividend for the high-demand problem (e.g., 48106) did not contain any multiples of 7. In Exp. 2, the difference between the low- and high-demand problems was whether or not the three faces shown on the 3D cube were adjacent to each other in the unfolded cube.

The flow of one trial in both experiments was described below (Figure 1A). There were 180 trials. At the start of each trial, participants choose either the left or right arrow. After the choice, an effort cue was presented. The effort cue was 100% associated with either low- or high-demand problems. After the effort cue, a problem was presented (high-demand or low-demand problem). The arrow determined the presentation probability of either low- or high-effort cues. The probabilities of arrows associated with upcoming effort cues, or effort levels, were alternated dynamically over time (Figure 1B). The set of either 80 or 20% was exchanged 5 times during 180 trials. After participants chose a cue with uncertainty, an effort cue was presented and participants knew which problem was to be presented with certainty. After that, a problem was presented, and they were asked to answer it. There was no time limit for response. Participants were instructed that their choices did not affect how fast they could finish the experiment.

In instruction, we did not explain to participants that there was a difference in the level of cognitive demands, between problem sets. Before the practice session(s), we gave verbal instructions and accepted questions from participants. After the practice session(s), participants were asked to explain what they thought the differences between the two types of problems were. If they answered that they thought the differences reflected differences in the actual content of the problems or differences in difficulty, we told them that we could not tell the answer was correct or not. If the difference reported by the participant was significantly wrong, we told them that the answer was wrong and asked them to practice. After the additional practice(s), if the participant subsequently reported things closer to reality, we told them that the answer was not incorrect. It should be noted that we never instructed the participants to avoid the high-demand problems to elicit voluntary choices. In addition, to minimize the influence of how fast the experiment could be completed, we instructed participants that their choices did not affect how fast they could finish the experiments.

Next, we outlined changes I–VII.

I: We added effort cues that corresponded exactly to the level of the upcoming effort to evoke cost PE signals. This is because the design of our previous experiments did not lock the time of cost PE generation. We used circle cues with red and green colors in Exp.

1. If participants were color blind, we planned to use blue instead of green but there were no color blindness participants. The color combinations between upcoming difficulties and colors were counterbalanced across participants. We used upward and downward triangle cues with grey color in Exp. 2. The combination between upcoming difficulties and directions (upward or downward triangle) were also counterbalanced across participants.

II: We instructed that there were two types of problems and they are red and green problems in Exp. 1 and upward and downward problems in Exp. 2. In our previous study, we instructed just that there were two types of problems.

III: We changed the number of trials and some conditions in the practice block(s), comparing with our previous work. Participants executed practice block(s) for 15-75 trials (mean ± standard deviation (SD), 26.5 ± 16.1) in Exp. 1 and 15-105 trials (mean ± SD, 32.1 ± 20.1) in Exp. 2. We did not give oral instructions during the practice block(s). Interstimulus intervals from the choice timing of arrows to the presentation timing of effort cues were 0.25, 0.50, 0.65 seconds jittered in Exp. 1 and 2.50, 5.00, 6.50 seconds jittered in Exp. 2. The duration of effort cue presentation was 0.50 seconds in both Exp. 1 and 2. Interstimulus intervals from the presentation timing of effort cues to the presentation timing of problems were 0.25 seconds in Exp. 1 and 2.50 seconds in Exp. 2. Intertrial intervals from the timing of answer for problems to the presentation timing of arrows of a next trial were 0.25, 0.50, 0.65 seconds jittered in Exp. 1 and 2.50, 5.00, 6.50 seconds jittered in Exp. 2. The mapping of arrows to demand levels were counterbalanced across, and stable in, both Exp. 1 and 2. After practice block(s), we asked participants how red and green problems in Exp. 1 and how upward and downward triangle problems differed to prevent misunderstanding the difference between the two kinds of problems (low- and high-demand problems). Notably, we did not explain the difference. If participants did not reply that the problems were low-versus high-demand, we responded that something else differentiated the problem types.

IV: Participants completed180 total trials in the scanner (the same number as our previous design, but we changed the number of blocks from four to six. More specifically, there were 6 blocks multiplied by 30 trials in both Exp. 1 and 2.

V: There were 180 trials totally, which consisted of 6 blocks of 30 trials, in both Exp. 1 and 2 in the scanner. In each block, a rest period was imposed until 200, 400, and 600s in Exp. 1 and 2 after the completion of the 10th, 20th, and 30th trial in the block in <200, 400, and 600s in both Exp. 1 and 2, respectively.

VI: We checked whether participants are right-handed or not with the FLANDERS handedness questionnaire ^45^.

VII: After the scan, participants answered questionnaires with some questions in both Exp. 1 and Exp. 2.

In Exp. 1, the first question was that “Did you feel that the following accurately describes the probabilities of red and green cues?”

After choosing an arrow, effort cues are presented that are 100% associated with the task types. The problems that followed the effort cues are always the division of 5-digit numbers, but there are two types of problems. The probability that each type of problem is more likely to appear is set independently for the left and right arrows. These probabilities changed very slowly over time. The same situation will continue for at least 10 trials. The presentation probabilities of which effort levels came after an arrow was chosen was maintained during the transition from block to block.

Participants then responded yes or no by circling their answer.

If the subject answered No, the participants answered the second question. “If you answered No on the above-mentioned question, please explain what was the difference specifically.”

The third question with a 7-point scale was that “How much did you like to solve either one of the red or green problems? Please draw a circle on the relevant vertical line on the horizontal bar below.”

In Exp. 2, we changed the description of the first question of Exp. 1 from the division of 5-digit numbers to the spatial reasoning problems and from red and green cues to upward or downward triangles. The second and third questions were the same.

### Image acquisition

We used a Phillips Ingenia 3.0T CX scanner with a 32-channel head coil (Shimane Institute of Health Science, Izumo, Shimane, Japan). Participants wore earplugs except for three participants of Exp. 1. Soft pads were used to reduce their head movements. The stimuli were presented via a mirror where the images of the monitor were reflected. Functional images were acquired in ascending order by T2-weighted echo planner imaging sequences [Flip angle: 80°; repetition time (TR): 2500ms; voxel size: 3.31mm × 3.37mm × 3.20mm; slice gap: no; field of view (FOV): 211.8 mm (Left-Right) × 212.3 mm (Anterior-Posterior) × 128.0 mm; slice number: 40; acquisition matrix: 64 × 63 × 40; volume number: 240 volumes × 6 blocks in both Exp. 1 and Exp. 2; parallel to the anterior and posterior commissures line]. We discarded the following 9 scans in each block to obtain the steady-state magnetization: the Phillips scanner deleted the first four scans automatically and we deleted the next five scans in each block for each participant.

### Behavioral Analysis

#### Basic Analysis

Behavioral analyses were the same protocols as our previous work (please see Mitsuto et al., 2018), except for the analyses of Generalized Linear Mixed Model (GLMM). We set the same inclusion criteria of correct rate with our previous work (> 80% for both high- and low-demand problems) and one participant was excluded in both experiments. We conducted a χ^2^ test and analyzed the effect of the demand that participants experienced two trials before.

#### Generalized Linear Mixed Model

We conducted GLMM analyses on the binominal choice data to test whether or not multiple past events affect current choices for demand avoiders who showed the avoidance behavior significantly in the aforementioned χ^2^ test. We use lmer4 package of R Studio for the GLMM analyses with maximum likelihood estimation methods. The response variable was coded as 1 for left choices and 0 for right choices. The fixed effects were an intercept, influences of choices and outcomes from both (k–1)-th to (k–4)-th trials’ choices on the k-th choices, and influences of choice repetitions. Specifically, the choice and outcomes were valued as 0 for both low-demand problems after left choices and high-demand problems after right choices and 1 for both high-demand problems after right choices and low-demand problems after left choices. The choice repetitions were valued as 0 for the opposite choices with the previous trials and 1 for the same choices as the previous trials. We used the z-score normalized values of them as the fixed effects. Additionally, the random effects were individual variations in the intercept, in the choices and outcomes of both (k–1)-th and (k–2)-th choices, and in choice repetitions.

The GLMM formulation was as follows:

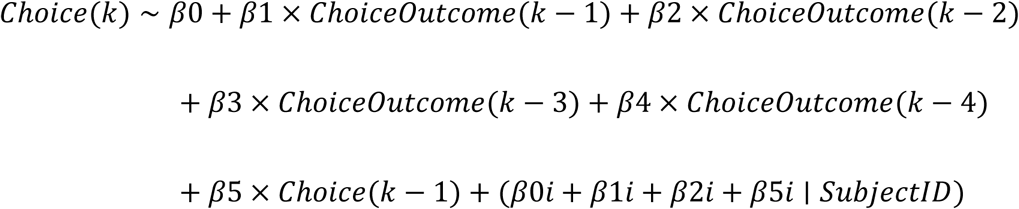

where β_0_ is the constant term, β_1_ to β_4_ are the effects of choice and outcomes from (k–1)- th to (k–4)-th trials on the choice at k-th trial, β_5_ is the effect of the choice at (k–1)-th trial, and β_0i_, β_1i_, β_2i_, and β_5i_ are the individual variations in these effects since individual differences in β1, β2, and β5 explained the most individual differences variance. Here, choices were binary ([Left, Right] = [0, 1]). The choice and outcome from (k–1)-th to (k– 4)-th trials ([Left_Low, Left_High, Right_Low, Right_High] = [0, 1, 1, 0]) and the choice at (k–1)-th trial ([Left, Right] = [0, 1]) were normalized in z-scores, respectively.

*Modelling, Model Fitting, and Comparisons.* The models that we used were the almost same set as our previous work (please see Mitsuto et al., 2018), except for the following change. We cut off the five worst-performing models, for which the expected cost for the unchosen option on the first trial was an additional free parameter. Concerning the following five reinforcement learning (RL) models, *ExpectedCost* for the unchosen option at the first trial [*ExpectedCost_unchosen_*(1)] was assumed to be equal to *ExpectedCost_Chosen_*(1).

1. solution time in individual trials (RL-Solution-Time-(ST) model)
2. demand level of the problem (RL-High-Low (HL) model)
3. incorrect solving (RL-Incorrect-Correct (IC) model)
4. sum of (1) and (3), with a weighting parameter for (3) (w_incorrect_) (RL-ST-IC model)
5. sum of (2) and (3), with a weighting parameter for (3) (w_incorrect_) (RL-HL-IC model) *ExpectedCost* for the unchosen and chosen options were assumed to be equal because means of the Bayesian information criterion (BIC) of the models where *ExpectedCost_unchosen_*(1) was set to be a free parameter were worse than the BIC means of the above-mentioned models that had the same initial values in our previous results and the presentation probabilities of which effort levels came after choices were 0.5 in practice sessions (please see the change III). Finally, we tested the following eleven models that were used in our previous work: a RL model based on solution time (RL-ST model), a low/high-demand model (RL-HL), a model based on incorrect responses (RL-IC),a model that combines solution time and incorrect responses ST + IC (RL-ST-IC), a model that combines low/high demands and incorrect choices (RL-HL-IC), a rest-time RL model (RL-Rest) and a rest-time model that also has information about low/high demands (RL-Rest-HL). Rest time model used a rest-time after the answer as a reinforcer in the reinforcement learning model. We also tested models based on Win-Stay-Lose-Shift probabilities (pWSLS), including information about low/high demands (pWSLS-HL), and incorrect responses (pWSLS-IC), and also a Probabilistic-Selection (PS) model based on high/low demands (PS-HL) and incorrect responses (PS-IC). The reason we used multiple RL models was to test whether the choice behavior was explained better by trial history. We used several probability models (WSLS and PS) because WSLS mode was widely-used models of choice behavior and PS model captured qualitative patterns in our data.

For convenience, we describe the RL-HL and PS-HL models below (which provided similarly good fits, according to model comparison, see Results), although the same description was also provided in our previous work ^11^. The RL-HL model was a RL model based on binarized effort cost, corresponding to low/high-demand. It was assumed that participants had the representation of the “Expected Cost (*ExpectedCost*)” of mental effort for options (i.e., arrows), this is, *ExpectedCost_left_*(*k*) and *ExpectedCost_right_*(*k*) (*k* = 1, 2, …, 180: trials). At each trial *k*, either the left or right arrow was assumed to be chosen with the probabilities *P_left_*(*k*) and *P_right_*(*k*), respectively:

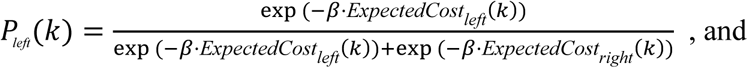

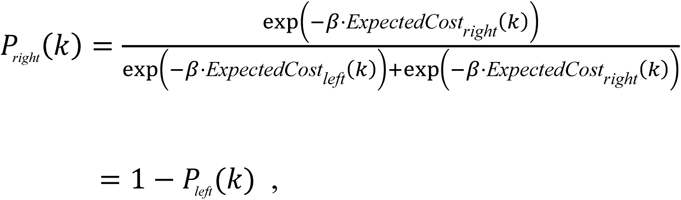

where *β* was a free parameter was the inverse temperature. The “Cost Prediction Error (*CPE*)” was also assumed to be calculated:

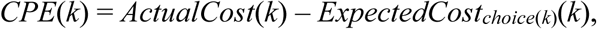

where Choice(*k*) was Left or Right depending on which was chosen. As *ActualCost*(*k*), we considered demand level of the problem, more specifically, 1 and 0 for high- and low-demand problems, respectively.

*ExpectedCost* for the chosen option was then assumed to be updated as follows:

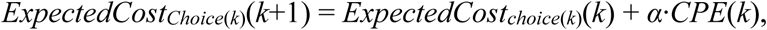

where *α* was a free parameter representing the learning rate. *ExpectedCost* for the unchosen option was assumed to be unchanged.

The PS-HL model was a PS model based on the experiences of low/high-demand. Participants repeated their choice from the preceding trial with the probability a and b after solving low- and high-demand problems, respectively.

Please note that we referred to the “RL model” in this article as the “PE model” in our previous article. As a result of model comparisons (see Results), we judged that it was reasonable to use the RL-HL model in the model-based fMRI analyses described below.

In sum, we analyzed: RL, rest-time RL, and probabilistic models; RL-ST model, RL-HL model, RL-IC model, RL-ST-IC model, RL-HL-IC model, RL-Rest model and RL-Rest-HL model, and probabilistic models including pWSLS-HL model and pWSLS-IC model, and PS-HL model and PS-IC model.

### Functional Imaging Analysis

We used SPM12 (Wellcome Trust Centre for Neuroimaging, UCL, London, https://www.fil.ion.ucl.ac.uk/spm/software/spm12/) for both preprocessing and analysis of functional imaging data. The preprocessing included the following steps: realigning to the first images using a six-parameter rigid body transformation, slice timing correction, coregistration, normalization, smoothing (with Gaussian kernel with a full-width at half-maximum of 8 mm). We set the inclusion criterion of both head movement (≤ 3mm) and behavioral performance (the above-mentioned high correct rate and the significant avoidance of high-demand problems on a χ^2^ test (p < 0.01)) as same as our previous work. We excluded 2 and 1 (Exp. 1 and 2, respectively) participants with >3 mm head movements among the participants who met the performance criterion and showed significant avoidance of high-demanding problems (see the Results). We defined striatum as dorsal striatum (DS, putamen and caudate) and VS (nucleus accumbens and the surrounding regions) in this article.

We fit general linear models (GLM) to our BOLD data (Exp. 1, n=25; Exp. 2, n=23). To turn off the default orthogonalization for multiple parametric modulations as same as our previous work, we commented out the lines 115-117 of spm_get_ons.m and the lines 235-245 of spm_fMRI_design.m. The analyses of both individual and group levels were done at the whole-brain in each experiment.

We used three GLMs (Figure 2) at the individual level analyses. In a trial, there were five events: arrow-cue presentation, the choice of an arrow, effort cue presentation, problem presentation, and the answer of problems.

First, we designed the GLM1 to test the first hypothesis that expected costs are updated at effort cues, by including a CPE regressor at the time of effort cues. Next, we designed GLM2 and GLM3 with the CPE regressor at the time of problem presentation or the when participants answered problems.

The trial-by-trial values of both *ExpectedCost* for the chosen option (referred to as *ExpectedCost_chosen_*) and the Cost Prediction Error (*CPE*) for individual participants were estimated by the RL-HL model. The response time from the arrow-cue presentation to the choice of an arrow was referred to as *RT_choice_*.

GLM1∼3 were designed in the following manner. We set the regressors at the time of arrow-cue presentation with parametric modulations by *ExpectedCost_chosen_* and *RT_choice_*, the regressors at the time of effort cue presentation with parametric modulations by problem-demand (0 and 1 for low- and high-demand problems, respectively) and CPE (this parametric modulation of CPE is only for GLM1), the regressor at the time of problem presentation with parametric modulation by CPE (this regressor with parametric modulation of CPE is only for GLM2), the regressors with the duration from problem presentation to the answer of problems with parametric modulations by problem-demand, the regressors at time of the answer of problems with parametric modulation by CPE (this regressor with parametric modulation of CPE is only for GLM3), the regressors for motor response at the time of the choice of an arrow and the answer of problems (GLM1 and GLM2 included them both timing and GLM3 included them only at the time of the choice of an arrow) and head movements. Each regressor in each GLM was convolved with the canonical hemodynamic response function, and one-sample t-tests were performed on individual maps of the regressors of interest for 25 participants in Exp. 1 and 23 participants in Exp. 2. The variance inflation factor (VIF) was calculated to determine whether the collinearity of the regressors of interest was within an acceptable range where the cut-off value for the VIF used in collinearity issues was 5 or 10.

For GLM 1-3, the variance inflation factors (VIF) of CPE parametric regressors at three timings were smaller than 5 across sessions and participants, which was judged it to be acceptable for collinearity.

Our conjunction analyses with a binary mask was the same as our previous work (please see Mitsuto *et al.*, 2018). We used the strict mask of common voxels between the results of Exp. 1 and Exp. 2 exceeding the threshold of cluster-level FWE corrected *p* < 0.05 and voxel-level uncorrected *p* < 0.001. We then reported correlates detected in the masked conjunction analyses with a threshold of family wise error (FWE) corrected p < 0.05 and voxel-level uncorrected *p* < 0.001. The relaxed mask reflected common voxels between Exp. 1 and Exp. 2 with the threshold of voxel-level uncorrected *p* < 0.01. We reported correlates detected in the relaxed masked conjunction analyses with a threshold of voxel-level uncorrected p < 0.001.

#### Residual Blood Oxygenation Level Dependent (BOLD) analyses

To test whether residual BOLD signals predict choice on each subsequent trial, we fit a version of our GLM1 above, but without a CPE regressor, and extracted residuals from each of the CPE-correlated cluster’s for each timing: at effort cue, effort initiation, and effort completion. CPE-correlated cluster regions of interest (ROIs) were the significant cluster of strict masked conjunction analyses with a threshold of FWE corrected p < 0.05 and voxel-level uncorrected p < 0.001: the right IPS from at effort cue, and the vmPFC/vACC/OFC from at effort initiation and the dmFC/dACC, left aINS, left M1/S1/IPS, and bilateral Cd from at effort completion.

We calculated the average residual BOLD signal across all voxels within each cluster at each time point, starting from time zero as the initial time point, for effort cue, effort initiation, and effort completion, up to 17.5 seconds. Subsequently, we averaged the residual BOLD signals between 5.0 and 7.5 seconds for each cluster, targeting the peak of canonical hemodynamic response function.

Next, we fit a generalized linear mixed effects model (with logit link function), regressing choice onto residual BOLD signal as follows:

𝐶ℎ𝑜𝑖𝑐𝑒(𝑘 + 1) ∼ 𝛽0 + 𝛽1 × 𝑅𝑒𝑠𝑖𝑑𝑢𝑎𝑙𝐵𝑂𝐿𝐷(𝑘) + (𝛽0𝑖 ∣ 𝑆𝑢𝑏𝑗𝑒𝑐𝑡𝐼𝐷)

where β_0_ is the constant term, β_1_ is the effects of residual BOLD signals at k-th trial, and β_0i_ is the individual variation in the constant term effects. Here, choices at (k + 1)-th trials were binary ([Stay, Switch] = [0, 1]). The residual BOLD signals at the k-th trial were normalized in two steps: first, within each block using z-scores, and second, across all blocks for each subject using z-scores.

As a sanity check, we also sought to confirm the relationship between trial-wise CPEs and trial-wise residual BOLD signals for each cluster at the time point, by performing Linear Mixed Model (LMM) analyses with the LMM was formulated as follows:

𝐶𝑃𝐸(𝑘) ∼ 𝛽0 + 𝛽1 × 𝑅𝑒𝑠𝑖𝑑𝑢𝑎𝑙𝐵𝑂𝐿𝐷(𝑘) + (𝛽0𝑖 + 𝑅𝑒𝑠𝑖𝑑𝑢𝑎𝑙𝐵𝑂𝐿𝐷(𝑘) ∣ 𝑆𝑢𝑏𝑗𝑒𝑐𝑡𝐼𝐷)

where β_0_ was the constant term, β_1_ was the effects of residual BOLD signals at k-th trial, and β_0i_ and β_1i_ were the individual variations in these effects. The residual BOLD signals at the k-th trial were normalized using the same two-step process described above, consistent with the normalization used in the GLMM.

Our mixed effects model analyses were conducted using MATLAB (version 2019b) with the Statistics and Machine Learning Toolbox, specifically using the *fitglme* and *fitlme* functions for GLMM and LMM, respectively.

Regarding missing data, trials with missing BOLD signals due to scan durations exceeding ten minutes per block were excluded from these fMRI analyses.

### Resource Availability

Upon request, the codes, the behavior data, and the preprocessed MRI data of this study are available from the corresponding author.

---

**Figure S1.**
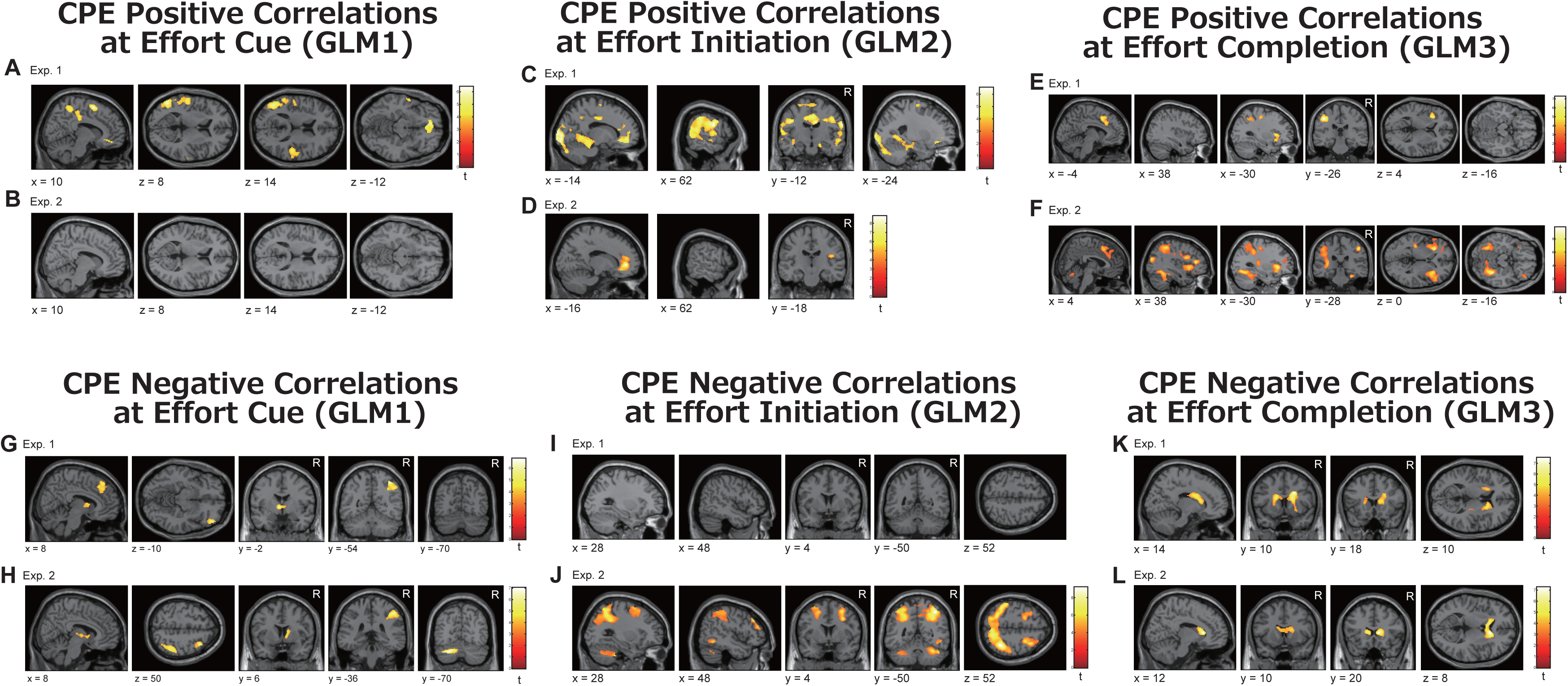
Neural correlates of the cost prediction error (CPE) at the time of effort cue, at effort initiation, and at effort completion. Panels A–F show positive correlations with CPE at the time of effort cue (GLM1), effort initiation (GLM2), and effort completion (GLM3) in Experiments 1 and 2. Panels G–L show corresponding negative correlations at the same stages. Results are thresholded at cluster-level FWE-corrected p < 0.05 or p ≈ 0.05 and voxel-wise uncorrected p < 0.001.

**Figure S2.**
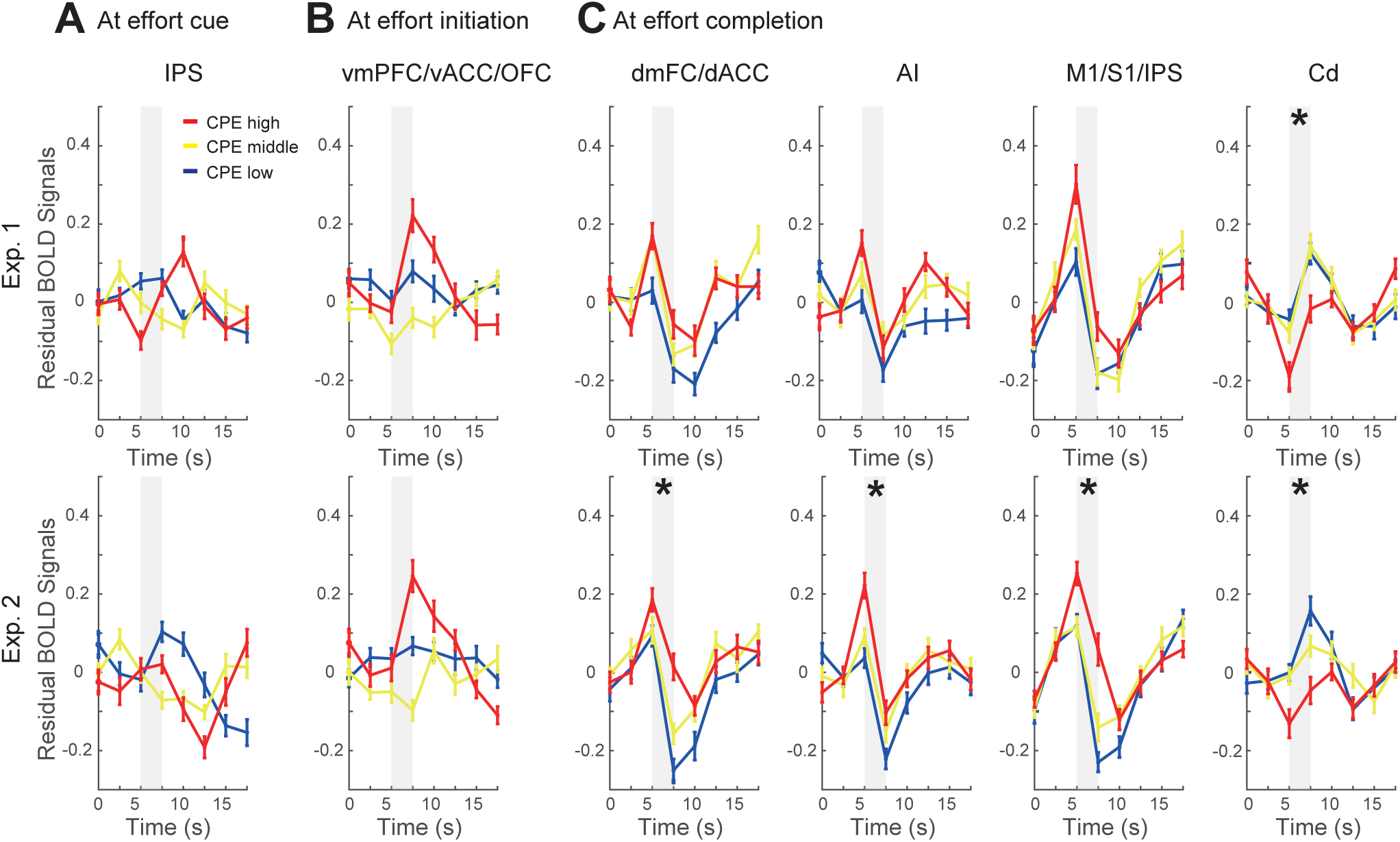
Residual BOLD signals predicted the sequential choices. Time-course analysis of residual BOLD signals for selected ROIs: the right IPS at effort cues, the vmPFC/vACC/OFC at effort initiation, and the dmFC/dACC, left aINS, left M1/S1/IPS, and bilateral Cd at effort completion. Curves represent residual BOLD signals averaged by CPE tertiles: CPE Low (−1 ≤ CPE < 33rd percentile, blue line), CPE Middle (33rd ≤ CPE ≤ 67th percentile, yellow line), and CPE High (67th percentile < CPE ≤ 1, red line). Before taking group averages, the residual BOLD signals for each participant were normalized in two steps: first, within each block using z-scores, and second, across all blocks for each subject using z-scores. Shaded regions indicate time points (5s and 7.5s) where the averaged values were used for analysis. Asterisks mark significant t-tests at p ≤ 0.05 in the GLMM. The bottom panels of Figure S2C are identical to those in Figure 7B.

**Table S1.**
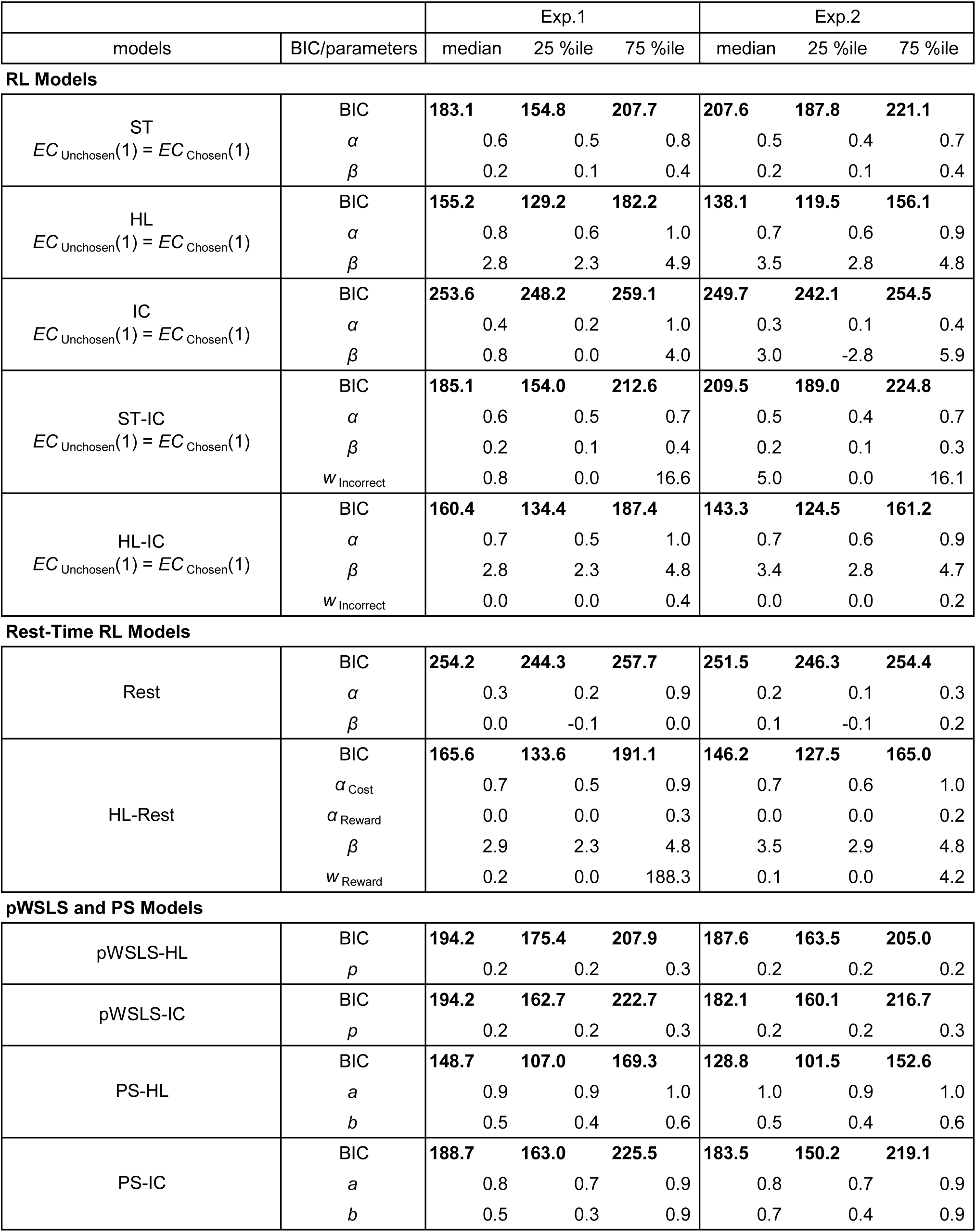
BIS scores and best-fit parameters for demand avoiders. This table shows the median, 25th and 75th percentiles of BIC scores and best-fit parameter estimates for 5 variants of reinforcement learning (RL) models with ActualCost (Solution-Time (SV), High-Low (HL), Incorrect-Correct (IC), ST-IC, and HL-IC), 2 variants of rest time considered RL models (Rest and HL-Rest models), and probabilistic Win-Stay-Lose-Shift (WSLS) and Probabilistic-Selection (PS) models. ECunchosen: the expected cost for the unchosen option, ECchosen: the expected cost for the chosen option, α: learning rate, β: inverse temperature, wincorrect: weighting parameter, αcost: the learning rate for cost, αreward: the learning rate for reward, wreward: weighting parameter, p: probability to select the different and same option after Win and Lose, respectively, a: probability to select the same option after Win, b: probability to select the same option after Lose.

**Table S2.** Neural correlates of cost-prediction-error at the time of effort cues using GLM1.

| Area | LR | cluster<br>size | cluster<br>p (FWE) | peak<br>T | MNI {mm} |  |  |
| --- | --- | --- | --- | --- | --- | --- | --- |
|  |  |  |  |  | x | y | z |
| **A Exp. 1** |  |  |  |  |  |  |  |
| *Positive correlation* |  |  |  |  |  |  |  |
| SMC/PCC | B | 3720 | <0.001 | 6.40 | 10 | 6 | 60 |
| MTG/AG | L | 2113 | <0.001 | 5.92 | -50 | -64 | 8 |
| vmPFC | B | 497 | <0.001 | 5.64 | -6 | 46 | -12 |
| PO | R | 426 | <0.001 | 4.83 | 50 | -24 | 14 |
| *Negative correlation* |  |  |  |  |  |  |  |
| IPS | R | 1095 | <0.001 | 6.80 | 38 | -54 | 54 |
| Thalamus | L | 289 | 0.038 | 6.10 | -6 | -2 | 2 |
| OFC/dIPFC | R | 974 | <0.001 | 5.84 | 50 | 44 | -10 |
| dmFC | B | 591 | 0.002 | 5.17 | 8 | 38 | 50 |
| **B Exp. 2** |  |  |  |  |  |  |  |
| *Positive correlation* |  |  |  |  |  |  |  |
| null |  |  |  |  |  |  |  |
| *Negative correlation* |  |  |  |  |  |  |  |
| Cerebellum | L | 568 | <0.001 | 7.02 | -22 | -70 | -30 |
| IPS/AG | R | 1439 | <0.001 | 6.35 | 48 | -36 | 50 |
| FEF/dIPFC | R | 720 | 0.001 | 6.31 | 36 | 18 | 54 |
| Caudate/TP | R | 338 | 0.034 | 4.89 | 12 | 6 | 10 |
| **C Conjunction analysis (strict mask <sup>(1)</sup>)** |  |  |  |  |  |  |  |
| *Positive correlation* |  |  |  |  |  |  |  |
| null |  |  |  |  |  |  |  |
| *Negative correlation* |  |  |  |  |  |  |  |
| IPS | R | 710 | 0.002 | 5.88 | 44 | -42 | 52 |
| **D Conjunction analysis (relaxed mask <sup>(2)</sup>)** |  |  |  |  |  |  |  |
| *Positive correlation <sup>(3)</sup>* |  |  |  |  |  |  |  |
| vmPFC | B | 114 | 0.365 | 4.10 | -4 | 44 | -12 |
| CO | R | 4 | 0.974 | 3.46 | 42 | 0 | 12 |
| *Negative correlation <sup>(3)</sup>* |  |  |  |  |  |  |  |
| IPS | R | 821 | <0.001 | 5.88 | 44 | -42 | 52 |
| OFC | R | 189 | 0.163 | 5.01 | 50 | 44 | -10 |
| FEF/dIPFC | R | 236 | 0.101 | 4.71 | 36 | 20 | 54 |

| Area | LR | cluster | cluster | peak | MNI {mm} |  |  |
| --- | --- | --- | --- | --- | --- | --- | --- |
|  |  | size | p (FWE) | T | x | y | z |
| Cerebellum | L | 137 | 0.285 | 4.56 | -30 | -68 | -30 |
| dIPFC | R | 65 | 0.610 | 4.35 | 48 | 40 | 20 |
| IC | R | 95 | 0.448 | 4.26 | 12 | 8 | 4 |
| dmFC | B | 267 | 0.074 | 4.14 | 6 | 30 | 46 |
| dIPFC | L | 57 | 0.659 | 3.90 | -42 | 52 | -2 |
| IPS | L | 24 | 0.871 | 3.90 | -44 | -48 | 52 |
| Cd | L | 15 | 0.923 | 3.90 | -8 | 8 | 4 |
| al | R | 49 | 0.711 | 3.72 | 32 | 20 | -6 |
| IFG | R | 11 | 0.943 | 3.46 | 54 | 20 | 8 |
| Cerebellum | L | 4 | 0.974 | 3.41 | -8 | -80 | -30 |
| Thalamus | R | 3 | 0.978 | 3.38 | 6 | -14 | 6 |

(1): a binary mask consisting of common voxels at cluster-level FWE corrected $p < 0.05$ or $p \approx 0.05$ and voxel-level uncorrected $p < 0.001$
(2): a binary mask consisting of common voxels at voxel-level uncorrected $p < 0.01$
(3): voxel-level uncorrected $p < 0.001$

**Table S3.** Neural correlates of cost-prediction-error at the time of effort initiation using GLM2.

| Area | LR | cluster<br>size | cluster<br>p (FWE) | peak<br>T | MNI {mm} |  |  |
| --- | --- | --- | --- | --- | --- | --- | --- |
|  |  |  |  |  | x | y | z |
| **A Exp. 1** |  |  |  |  |  |  |  |
| *Positive correlation* |  |  |  |  |  |  |  |
| S1/V2/EC/FC | B | 17659 | <0.001 | 6.52 | 62 | -6 | 28 |
| S1/PC | L | 2544 | <0.001 | 5.54 | -44 | -12 | 32 |
| SMC/PMC | B | 262 | 0.045 | 4.56 | -24 | -10 | 68 |
| *Negative correlation* |  |  |  |  |  |  |  |
| null |  |  |  |  |  |  |  |
| **B Exp. 2** |  |  |  |  |  |  |  |
| *Positive correlation* |  |  |  |  |  |  |  |
| vmPFC/vACC | B | 2986 | <0.001 | 8.77 | 16 | 44 | 2 |
| Operculum | R | 489 | 0.008 | 5.95 | 42 | -18 | 20 |
| *Negative correlation* |  |  |  |  |  |  |  |
| null |  |  |  |  |  |  |  |
| **C Conjunction analysis (strict mask <sup>(1)</sup>)** |  |  |  |  |  |  |  |
| *Positive correlation* |  |  |  |  |  |  |  |
| vmPFC/vACC/OFC | B | 961 | <0.001 | 5.37 | -14 | 34 | -8 |
| *Negative correlation* |  |  |  |  |  |  |  |
| null |  |  |  |  |  |  |  |
| **D Conjunction analysis (relaxed mask <sup>(2)</sup>)** |  |  |  |  |  |  |  |
| *Positive correlation <sup>(3)</sup>* |  |  |  |  |  |  |  |
| vmPFC/vACC/OFC | B | 1522 | <0.001 | 5.37 | -14 | 34 | -8 |
| OFC | R | 9 | 0.955 | 4.31 | 28 | 34 | -12 |
| dACC | L | 30 | 0.836 | 3.99 | -2 | 30 | 16 |
| PO | R | 47 | 0.724 | 3.91 | 46 | -22 | 20 |
| CO | R | 17 | 0.914 | 3.43 | 48 | -10 | 16 |
| STC | R | 3 | 0.979 | 3.41 | 66 | -40 | 20 |
| dmPFC | R | 7 | 0.964 | 3.40 | 2 | 52 | 18 |
| MTC | R | 4 | 0.976 | 3.33 | 46 | -38 | 4 |
| MTC | L | 1 | 0.987 | 3.32 | -60 | -2 | -18 |
| MTC | L | 1 | 0.987 | 3.30 | -58 | 0 | -20 |
| *Negative correlation <sup>(3)</sup>* |  |  |  |  |  |  |  |
| null |  |  |  |  |  |  |  |

(1): a binary mask consisting of common voxels at cluster-level FWE corrected $p < 0.05$ or $p \approx 0.05$ and voxel-level uncorrected $p < 0.001$
(2): a binary mask consisting of common voxels at voxel-level uncorrected $p < 0.01$
(3): voxel-level uncorrected $p < 0.001$

**Table S4.** Neural correlates of cost-prediction-error at the time of effort completion using GLM3.

| Area | LR | cluster<br>size | cluster<br>p (FWE) | peak<br>T | MNI {mm} |  |  |
| --- | --- | --- | --- | --- | --- | --- | --- |
|  |  |  |  |  | x | y | z |
| **A Exp. 1** |  |  |  |  |  |  |  |
| *Positive correlation* |  |  |  |  |  |  |  |
| S1/M1 | L | 1397 | <0.001 | 7.25 | -44 | -26 | 46 |
| al | L | 368 | 0.012 | 5.75 | -32 | 18 | 4 |
| dmFC/dACC | B | 635 | <0.001 | 5.49 | -6 | 14 | 46 |
| *Negative correlation* |  |  |  |  |  |  |  |
| MFG/Cd | B | 2212 | 0.000 | 7.51 | 24 | 10 | 30 |
| **B Exp. 2** |  |  |  |  |  |  |  |
| *Positive correlation* |  |  |  |  |  |  |  |
| FG/Cerebellum | R | 1899 | <0.001 | 9.74 | 34 | -48 | -16 |
| al | L | 1005 | <0.001 | 8.18 | -28 | 24 | 0 |
| IPS/MOG | R | 3319 | <0.001 | 8.00 | 48 | -28 | 48 |
| dIPFC/AI | R | 2323 | <0.001 | 7.45 | 38 | 36 | 14 |
| FG/Cerebellum | L | 953 | <0.001 | 6.88 | -30 | -70 | -12 |
| PI/IPS/M1 | L | 3103 | <0.001 | 6.29 | -36 | -24 | -2 |
| dmFC/dACC | B | 1346 | <0.001 | 5.87 | 4 | 18 | 46 |
| IPS | L | 384 | 0.011 | 5.10 | -22 | -70 | 32 |
| MFG/SFG | L | 289 | 0.033 | 5.00 | -22 | 46 | 10 |
| IFG/PCG | L | 340 | 0.018 | 4.99 | -56 | 12 | 24 |
| *Negative correlation* |  |  |  |  |  |  |  |
| Cd | B | 1062 | <0.001 | 7.24 | 12 | 20 | 8 |
| **C Conjunction analysis (strict mask <sup>(1)</sup>)** |  |  |  |  |  |  |  |
| *Positive correlation* |  |  |  |  |  |  |  |
| M1/S1/IPS | L | 940 | <0.001 | 6.01 | -44 | -26 | 46 |
| al | L | 322 | 0.032 | 5.62 | -30 | 22 | 2 |
| dmFC/dACC | B | 423 | 0.012 | 5.07 | -4 | 14 | 50 |
| *Negative correlation* |  |  |  |  |  |  |  |
| Cd | B | 371 | 0.019 | 5.42 | 14 | 18 | 10 |
| **D Conjunction analysis (relaxed mask <sup>(2)</sup>)** |  |  |  |  |  |  |  |
| *Positive correlation <sup>(3)</sup>* |  |  |  |  |  |  |  |
| S1/M1/IPS | L | 1092 | <0.001 | 6.01 | -44 | -26 | 46 |

| Area | LR | cluster | cluster | peak | MNI {mm} |  |  |
| --- | --- | --- | --- | --- | --- | --- | --- |
|  |  | size | p (FWE) | T | x | y | z |
| al | L | 434 | 0.011 | 5.62 | -30 | 22 | 2 |
| dmFC/dACC | B | 644 | 0.002 | 5.07 | -4 | 14 | 50 |
| FG | L | 78 | 0.518 | 4.72 | -30 | -72 | -10 |
| AL/FO | R | 283 | 0.048 | 4.52 | 36 | 22 | 4 |
| FG | R | 220 | 0.097 | 4.47 | 34 | -48 | -20 |
| dIPFC | R | 60 | 0.633 | 4.26 | 50 | 34 | 16 |
| PCG/MFG | L | 29 | 0.850 | 4.12 | -46 | 6 | 32 |
| PCG/IFG | R | 105 | 0.377 | 4.01 | 46 | 8 | 26 |
| SMG | R | 39 | 0.781 | 3.93 | 46 | -40 | 42 |
| Precuneus | R | 13 | 0.945 | 3.91 | 20 | -62 | 26 |
| OFC | R | 21 | 0.901 | 3.68 | 32 | 18 | -20 |
| MFG | L | 10 | 0.959 | 3.56 | -28 | 4 | 44 |
| AG | R | 9 | 0.963 | 3.49 | 56 | -48 | 28 |
| FG | R | 3 | 0.985 | 3.49 | 32 | -34 | -22 |
| FG | R | 1 | 0.991 | 3.37 | 32 | -30 | -24 |
| MFG | L | 2 | 0.988 | 3.34 | -22 | 48 | 12 |
| MTG | L | 3 | 0.985 | 3.32 | -60 | -28 | -14 |
| SMG | R | 1 | 0.991 | 3.31 | 50 | -26 | 48 |
| *Negative correlation <sup>(3)</sup>* |  |  |  |  |  |  |  |
| Cd | B | 553 | 0.004 | 5.42 | 14 | 18 | 10 |
| Thalamus | B | 28 | 0.857 | 4.18 | 0 | -10 | 6 |
| Cd | L | 5 | 0.979 | 3.42 | -16 | 28 | 6 |
| Pallidum | L | 1 | 0.991 | 3.40 | -20 | -6 | -10 |

(1): a binary mask consisting of common voxels at cluster-level FWE corrected $p < 0.05$ or $p \approx 0.05$ and voxel-level uncorrected $p < 0.001$
(2): a binary mask consisting of common voxels at voxel-level uncorrected $p < 0.01$
(3): voxel-level uncorrected $p < 0.001$

